# Distinct proteostasis states drive pharmacologic chaperone susceptibility for Cystic Fibrosis Transmembrane Conductance Regulator misfolding mutants

**DOI:** 10.1101/2021.09.09.459524

**Authors:** Eli Fritz McDonald, Carleen Mae P. Sabusap, Minsoo Kim, Lars Plate

## Abstract

Pharmacological chaperones represent a class of therapeutic compounds for treating protein misfolding diseases. One of the most prominent examples is the FDA-approved pharmacological chaperone lumacaftor (VX-809), which has transformed cystic fibrosis (CF) therapy. CF is a fatal disease caused by mutations in the cystic fibrosis transmembrane conductance regulator (CFTR). VX-809 corrects folding of F508del CFTR, the most common patient mutation, yet F508del exhibits only mild VX-809 response. In contrast, rarer mutations P67L and L206W are *hyper-responsive* to VX-809, while G85E is *non-responsive*. Despite the clinical success of VX-809, the mechanistic origin for the distinct susceptibility of mutants remains unclear. Here, we use interactomics to characterize the impact of VX-809 on proteostasis interactions of P67L and L206W and compare these to F508del and G85E. We determine *hyper-responsive* mutations P67L and L206W exhibit decreased interactions with proteasomal, and autophagy degradation machinery compared to F508del and G85E. We then show inhibiting the proteasome attenuates P67L and L206W VX-809 response, and inhibiting the lysosome attenuates F508del VX-809 response. Our data suggests a previously unidentified but required role for protein degradation in VX-809 correction. Furthermore, we present an approach for identifying proteostasis characteristics of mutant-specific therapeutic response to pharmacological chaperones.

## INTRODUCTION

The proteostasis network (PN) evaluates the folding fidelity of proteins, allowing trafficking of properly folded proteins to their cellular destination while coordinating degradation of misfolded proteins. Misfolded proteins can be rescued from PN degradation by pharmacological chaperones, an emerging class of small molecule therapeutics that stabilize protein targets. For instance, the pharmacological chaperone Lumacaftor (VX-809) is part of an FDA-approved treatment for Cystic Fibrosis (CF), one of the most common lethal Mendelian diseases in the U.S. CF arises from mutations in the cystic fibrosis transmembrane conductance regulator (CFTR)^1,2,3^. CFTR regulates epithelial ion homeostasis necessary for many organ-liquid interfaces and CF patients suffer from altered mucus secretion leading to chronic lung infection, pancreatic and hepatic insufficiency, and premature death^4^. Current CF treatment involves stabilizing mutant CFTR with VX-809^1^ to restore protein levels by modulating folding and trafficking. However, VX-809 availability is limited to patients with a small number of mutations that are predicted to have corrector response based on *in vitro* evaluations typically prioritized based on patient population size^5–11^.

Among patients with mutations approved for VX-809 treatment, different CFTR mutations exhibit highly variable clinical efficacy^11^. For example, initial CFTR modulators were developed for the most common mutation F508del^12^. F508del represents the prototypical class-II disease-causing mutation leading to altered proteostasis^13^, trafficking,^14^ and decreased cell surface expression^15^. However, F508del exhibits mild VX-809 rescue relative to other class-II mutations, both *in vitro*^16^ and among clinical studies^17^. By contrast, the rare mutation P67L, which occurs in less than 300 patients^18^, demonstrates robust VX-809 correction with P67L reaching levels of mature CFTR at the cell surface comparable to wild type (WT) in some cell model systems^19^. Likewise, the mutation L206W responds well to VX-809 *in vitro*^20^. Still other mutations, such as G85E, appear to be completely resistant to VX-809 correction *in vitro*^21^. Despite the identical classification of F508del, P67L, L206W, and G85E variants as class-II (abnormal protein folding/trafficking)^22^, this classification scheme provides little basis for predicting CFTR therapeutic response, otherwise known as *theratype*^23,24^. F508del and G85E show similar theratypes with limited to no change in protein levels in response to VX-809 (*hypo-responders*) whereas P67L and L206W exhibit dramatic increases in protein levels in response to VX-809 (*hyper-responders*). We sought to understand the underlying proteostasis interactions contributing to CFTR mutant theratype, towards the goal of predicting pharmacological chaperone clinical efficacy given a patient mutation.

Previous studies attributed variable VX-809 response to the fundamental folding and/or structural defects conferred by any given mutation^25^. However, precisely how changes in structural defects translate to changes in protein expression levels and trafficking efficiency remains unclear. Folding and trafficking of CFTR variants is mediated by changes to quality control processes governed by chaperones, folding, trafficking, and degradation factors comprising the proteostasis network (PN). On the one hand, distinct mutations may cause distinct domain level conformational changes leading to changes in chaperone and degradation factor binding affinity^26,27^. On the other hand, different mutations may experience distinct rate-limiting steps in protein quality control. CFTR protein quality control is dictated by the CFTR-PN protein interactions that regulate the balance between trafficking and degradation^13,28^. Thus, each mutation may get trapped in a specific subcellular locale where the confluence of PN interactors leads to degradation^29–31^. Consequently, it is imperative to understand how VX-809 shapes mutation specific PN interactions^32,33^. A better mechanistic understanding of this interplay could allow CFTR variant theratyping for future corrector compounds and reveal other potential protein targets for CF treatment.

Past CFTR PN characterization focused on F508del under temperature correction or VX-809 treatment conditions^28,34^. Temperature correction to 30 °C modulates distinct F508del interactions with Hsp70 and Hsp90 chaperones, as well as several degradation factors to enable enhanced trafficking^29^. VX-809 treatment also modulates PN interactions and remodels the F508del interactome towards WT^35^. These studies identified chaperones as viable targets for altering CFTR trafficking, demonstrated the broad impact of VX-809 altered proteostasis, but offered little mechanistic insight into the function of VX-809. Few studies have investigated other CFTR mutations, particularly within the context of mutation-specific theratype.

Here, we used affinity purification-mass spectrometry (AP-MS) to sensitively detect and quantify CFTR interacting proteins and identify cellular machinery responsible for managing CFTR trafficking or degradation after VX-809 treatment. We quantitatively compared multiple mutants with DMSO and VX-809 treatment by multiplexing AP-MS with isobaric tandem mass tags (TMT)^36^. This approach allowed us to sensitively determine interactome changes and PN remodeling under VX-809 treatment for *hyper-responders* P67L and L206W CFTR. We found that *hyper-responsive* mutants shared similar proteostasis interactions to F508del with DMSO – consistent with their common class-II variant classification. However, both *hyper-responsive* mutants showed decreased interactions with proteasomal proteins and autophagy proteins under VX-809 treatment compared to F508del. We then showed that inhibition of the proteasome substantially attenuates the VX-809 response of *hyper-responsive* mutations P67L and L206W as well as partially attenuates the VX-809 response of F508del. Interestingly, inhibition of lysosomal degradation also attenuates F508del response, but not P67L or L206W. Together these data suggest a previously unidentified role for protein degradation in the VX-809 mechanism. Our multiplexed approach allows for parallel theratyping of additional CFTR variants with other CFTR correctors including recently FDA approved VX-445^37^. Furthermore, our methods can be applied to theratyping other protein misfolding diseases to emerging pharmacological chaperone therapies.

## RESULTS

### Co-immunoprecipitation of VX-809 hyper- and hypo-responsive CFTR variants

We sought to determine how the misfolding CFTR mutants differ in proteostasis interactomes from those of WT. Specifically, we focused on comparing VX-809 *hyper-responders* (P67L, L206W) to *hypo-responders* (F508del, G85E) (**Figure 1A**) to elucidate the mechanisms underlying the divergent VX-809 response. We used co-immunoprecipitation (Co-IP) coupled to LC-MS/MS analysis to isolate CFTR and identify its interacting proteins (**Figure 1B-C**)^38^. To enable direct, relative comparison of interactor abundance across multiple CFTR variants and in response to corrector treatment, we employed tandem mass tag (TMT) multiplexing^36^. Immunoblots confirmed the successful isolation of all four CFTR variants and relative amounts of band B and C were consistent with the protein levels observed in lysates (**Figure 1B**). As expected, we observed lower band C amounts for all the mutant variants (F508del, P67L, L206W, and G85E) compared to WT consistent with their protein trafficking defect. Importantly, we saw near restoration of band C levels for P67L and L206W with VX-809 treatment (*hyper-responders*), while F508del and G85E showed minimal to no band C in the presence of VX-809 (*hypo-responders*) (**Figure 1B**). After cleanup and proteolytic digestion, we then labeled different Co-IP conditions with respective TMT 11-plex reagents (WT and 4 mutants, each +/- VX-809, and mock transfection). We included a tdTomato mock transfection control to account for non-specific background during the Co-IP to distinguish CFTR-specific interactors. After TMT labeling, combinations of 11 distinct Co-IP conditions were pooled into a single mass spectrometry sample and analyzed by multidimensional protein identification technology (MudPIT)-tandem mass spectrometry. In addition to peptide identification, TMT reporter ions produced during peptide fragmentation enable direct, relative comparison of peptide abundances across the different Co-IP conditions (**Figure 1D**). We conducted Co-IPs and independent mass spectrometry runs for multiple replicates per mutation (WT N=10, P67L N= 12, F508del N=11, G85E N=7, L206W N=7) (**Supplemental Dataset 1**). We used these data to validate the TMT labeling scheme applicability to CFTR interactomics and to quantitively compare interactomes of *hyper*- and *hypo-responsive* mutants in response to VX-809.

**Figure 1.**
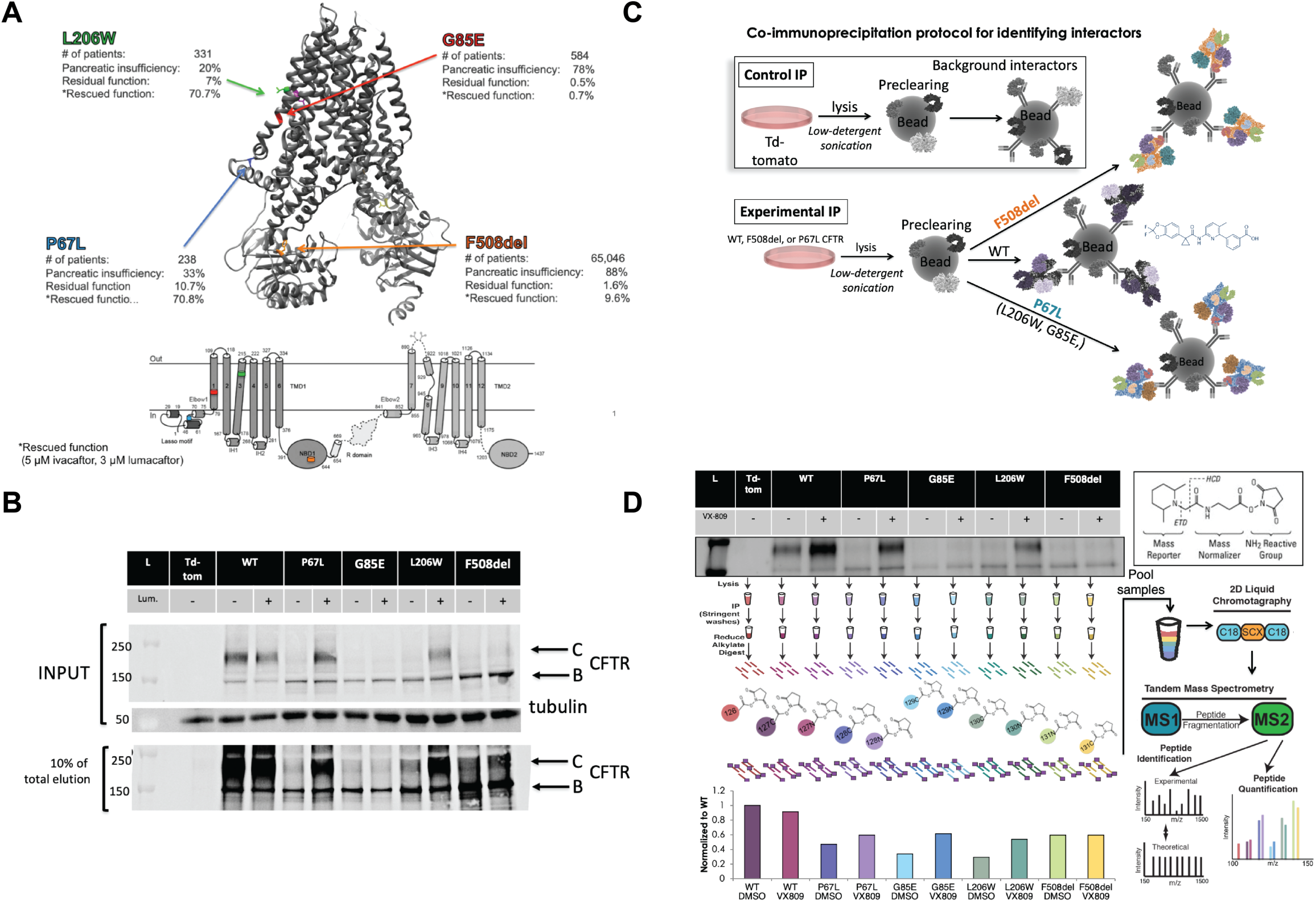
Multiplexed Affinity-Purification Mass Spectrometry (AP-MS) for identification of variant specific CFTR interactomes in response to VX-809. **A**. Schematic showing CFTR cryo-EM structure (PDB ID: 5UAK) and location of pathological mutations and annotation of mutations by CFTR domain. Disease severity and CFTR function in response to modulators are indicated^18,73^. **B**. Western blots showing expression and IP of CFTR variants in the absence or presence of 3µM VX-809. Input from total cell lysate is shown on top. Samples were pre-cleared with Protein G beads inputs purified on beads complexed to 24-1 CFTR antibodies. IP elution are shown at the bottom Identification of non-specific interactors used mock (non-CFTR expressing) cells^38^. **C**. Schematic of the IP procedure highlighting use of mock control and CFTR variants. Interactors likely change depending on the VX-809 response of each variant. **D**. Schematic of TMT-quantification AP-MS interactomics workflow and example of multiplexed analysis of mutations and treatments to measure interactomics changes. Duplicates of P67L and F508del treated samples with tdTomato (mock), and WT controls were purified via IP. Samples are digested, peptides are labeled with individual TMT reagents, and pooled for tandem mass spec analysis. Relative mass spec quantification of CFTR levels recapitulates biochemical detection via Western blot.

### Validation of TMT-based AP-MS proteomics to confirm WT and F508del CFTR interactors

Prior CFTR interactomics approaches have relied on co-interacting protein identification technology (CoPIT) with label-free quantifications^28,35,39^. To validate our TMT based quantification approach, we compared our identified interactions to previous WT and F508del CFTR studies. We determined statistically significant interactors by comparing the log2 TMT intensities of identified proteins in the CFTR co-IP samples to tdTomato mock transfected samples (**Figure 2A, Supplemental Figure S1**). In line with prior studies, we used median normalized TMT intensity for this comparison based on the assumption that the majority of identified proteins represent non-specific interactors that are present in similar amounts in the mock and CFTR-bait Co-IP samples (**Supplemental Figure S1A**, see materials and methods)^40,41^. WT and F508del CFTR demonstrated significant enrichment compared to tdTomato control (**Supplemental Figure S1B-E**). We defined statistically significant interactors as proteins using a combined p-value cutoff of 0.05 and a positive log2 fold change compared to tdTomato control. Our initial goal was to prioritize a comprehensive list of interactors that showed significant enrichment with any of the variants in our datasets (WT, F508del, P67L, or L206W). We compiled a list of enriched proteins for each of the individual comparisons to mock into a master list for quantitative comparison (**Figure 2A, Supplemental Dataset 2**). For validation, we compared the variant-specific interactor lists and the master list to prior CFTR interactome datasets.

**Figure 2.**
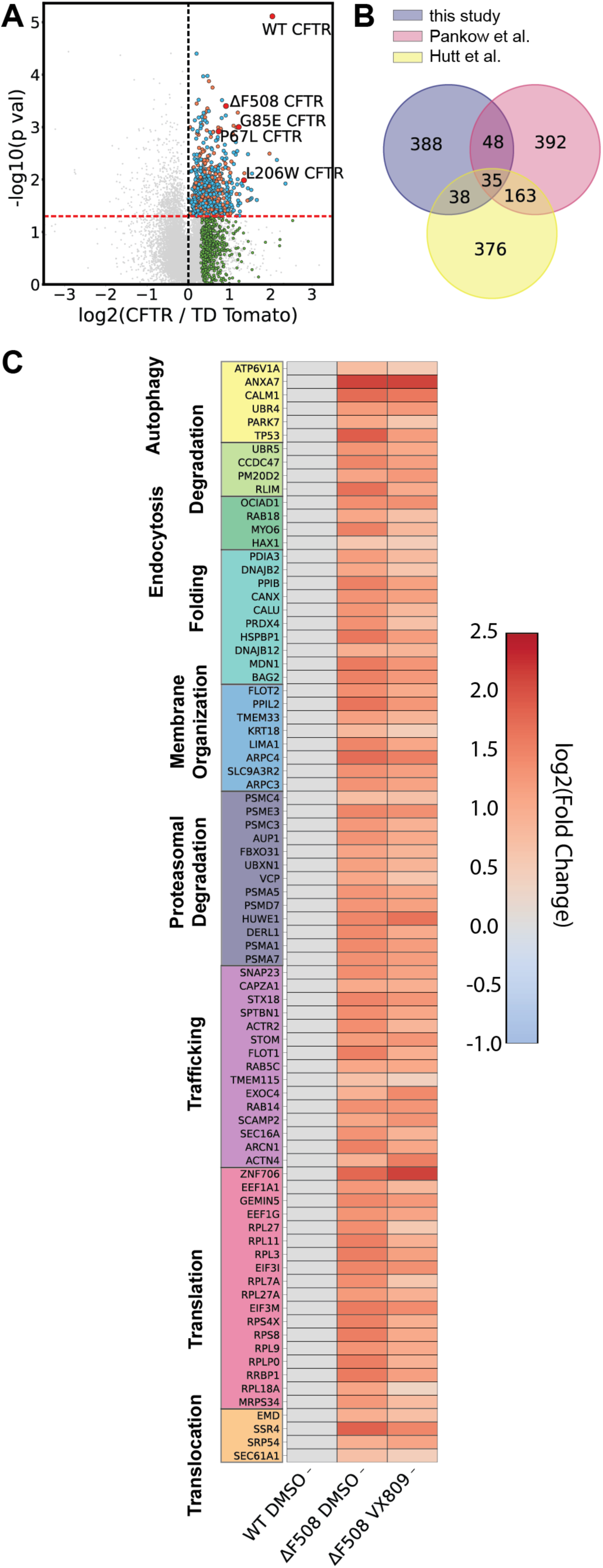
Identification and validation of altered F508del interactome with VX-809. **A**. Volcano plotting showing identification of CFTR variant interactors using multiplexed AP-MS proteomics. The x axis represents log 2-fold change of CFTR IP over IP of tdTomato mock control. A statistical significance cutoff of p > 0.05 is portrayed by an orange line. Blue dots represent newly identified statistically significant interactors in this study, orange dots represent overlap with previously identified interactors from Pankow, et al.^28^ and Hutt, et al.^35^ that meet our statistical threshold for interactors, and green dots represent previously identified interactors^28,35^ that did not meet our statistical threshold, but were enriched greater than one standard deviation of the log 2-fold change distribution between CFTR over tdTomato control. **B**. Overlap between statistically significant interactors identified in this study across variants considered with previously identified interactors from Pankow, et al. and Hutt, et al.^28,35^ **C**. Heatmap displaying interactions changes experienced by F508del CFTR under 3 μM VX-809 treatment compared to DMSO treatment. The log2 fold changes are scaled to WT with DMSO. Proteins were selected from manually curated list of proteostasis interactors and pathways were assigned with GO-terms.

We identified a total of 501 CFTR interactors with WT, F508del, P67L, L206W, and G85E CFTR with DMSO and VX-809 treatment conditions. Our CFTR interactome shares 93 and 73 interactors with prior datasets published by Pankow, et al. and Hutt, et al. respectively (**Figure 2B**)^28,35^. It is noteworthy that the overlap between prior datasets from Pankow, et al. and Hutt, et al. is relatively low (only ∼30%). Our degree of overlap with both datasets is comparable with ∼20% of identified interactors found in either study.

Nonetheless, we identify slightly fewer interactors in total compared to label-free proteomics studies. To understand the limitations of our approach, we also mapped previously identified interactors on volcano plots (shown in green in **Figure 2A, Supplemental Figure S1B-E)**. Several prior interactors were enriched in the CFTR-bait samples compared to the mock but did not pass our statistical confidence cutoffs. We also observed other prior interactors that were enriched inconsistently in our samples. Overall, the discrepancies may result from differences in the quantification approach and higher variance in our multi-step TMT sample processing procedure. At the same time, we cannot exclude the possibility that the prior label-free datasets suffered from a high false-positive discovery rate, which would explain the large number of interactors only observed in a single dataset (**Figure 2B**). Regardless, it is important to highlight that we consistently identified core proteostasis interactors with known function in CFTR biogenesis and trafficking (e.g. calnexin (CANX), SEC61A1, and MYO6) (**Supplemental Table 1**), validating our TMT-based quantitative AP-MS approach to characterize the CFTR proteostasis interactome.

We next used our master list of CFTR interactors (**Supplemental Dataset 2**) for subsequent quantitative comparison between CFTR variants and drug treatment conditions. We also included proteins that Pankow, et al. and Hutt, et al. previously identified as significant CFTR interactors that we identified as enriched greater than one standard deviation of log2 fold change distribution relative to our mock control (green dots in **Figure 2A, Supplemental Figure S1**). We categorized our grouped list of interactors by biological pathways and refined the classifications manually using the GO-terms for biological processes (see materials and methods) (**Figure 2C, Supplemental Dataset 2**). This analysis further highlighted that our TMT-based quantitative interactomics analysis is able to recapitulate the broad categories of CFTR proteostasis pathways identified in previous studies^28,35^.

### Quantitative comparison of F508del CFTR to confirm increased PN interactions

To further validate of our TMT approach, we compared interaction levels between WT and F508del in the presence and absence of VX-809. Importantly, direct quantitative comparison requires normalization to the bait protein (CFTR) to account for the inherently lower CFTR levels in F508del co-IP samples compared to WT (**Figure 1C, Supplemental Figure S2**, see materials and methods). We scaled the log2 fold abundances to WT DMSO condition within each run to provide a common reference point and plotted the resulting normalized interaction abundances as a heatmap organized by proteostasis pathways. We included autophagy, non-specific degradation, endocytosis, folding, membrane organization, proteasomal degradation, trafficking, translation, and translocation (**Figure 2C**) as proteostasis pathways of interest. Consistent with previous findings, this heatmap reveals an overall increase in proteostasis interactions with F508del compared to WT.

F508del interacts more than WT with previously characterized proteostasis factors, such as PARK7, BAG2, DNAJB12, HSPBP1, DNAJB2, Derlin1 (DERL1), and Calnexin (CANX) (**Figure 2C**). Previous studies showed F508del rescue through knockdown of PARK7^35^, BAG2^42^, DNAJB12^43^, and DERL1^44^ or decreasing CFTR levels with co-expression of DNAJB2^45^, indicating a pro-degradation role for these proteostasis factors. By contrast, other studies show F508del rescue through overexpression of Calnexin (CANX)^46^ and HSPBP1^47^, highlighting their pro-folding role. Consequently, our TMT quantification approach can recapitulate the enhanced interactions with these well characterized F508del interactors. We also find that F508del interacts more with trafficking machinery (**Figure 2C**), which is consistent with previous studies^28^. This could reflect increased F508del dwell time with ER trafficking machinery as the protein is unable to fully traffick to the plasma membrane. Finally, F508del interacts more strongly with translation and translocation machinery (**Figure 2C**), likely due to increased F508del retention in the ER, slower translation^48^, and membrane insertion^49^.

Next, we examined how VX-809 treatment alters the proteostasis interactions for F508del. The scaled log2 TMT abundances in our multiplexed experimental setup allow for direct comparison of interaction intensities (**Figure 2C**). Overall, VX-809 mildly reduces interaction with proteostasis factors compared to untreated F508del. However, interactions are still broadly increased relative to WT CFTR (**Figure 2C**). This observation is consistent with prior interactomics characterization of the F508del response to VX-809 by Hutt et al.^35^, and it is likely reflective of the incomplete rescue of F508del trafficking by the corrector drug. Furthermore, Pankow et al. observed reduced F508del interactions with proteostasis factors upon shift to low temperature (30 °C), which was able to restore mutant CFTR trafficking^28^. Similarly, we observed that VX-809 slightly reduces F508del interactions with folding machinery. Lastly, we saw some attenuation in interactions with known proteostasis factors that limit F508del trafficking, for example, BAG2 and PARK7^35,42^. Thus, VX-809 reduction of F508del interaction with BAG2 and PARK7 is consistent with VX-809 correcting F508del folding sufficiently to attenuate these pro-degradation interactions. We also examined WT CFTR interaction changes in the presence of VX-809 and found that several proteostasis interactions are reduced (**Supplemental Figure S3A**). The overall identification of these important proteostasis components and quantification of their down-regulation with VX-809 supports our ability to measure subtle protein interaction changes with TMT-labeling. With our approach validated, we then examined protein interaction changes for the *hyper-* and *hypo-responsive* CFTR variants.

### Class-II misfolding mutation P67L exhibits similar interactome to F508del

We first examined the class-II variant P67L, which is much more responsive to VX-809 treatment than F508del, exhibiting restoration of band C CFTR levels similar to WT (**Figure 1B**). The P67L mutation occurs in the lasso-motif helix-helix bend at the CFTR N-terminus (**Figure 1A**). The mutation prevents proper N-terminal/C-terminal interactions resulting in ER-retention and premature degradation^50^. Previous investigation of P67L by Western blot trafficking assay, Ussing chamber electrophysiology, sequence conservation analysis, and molecular dynamics simulation characterized P67L as a VX-809 *hyper-responder*^19,50^. However, the cellular proteostasis interactome of P67L remains unknown.

Since P67L and F508del share a similar fate in the ER as trafficking-defective mutants, we hypothesized P67L exhibits similar protein interactions to F508del compared to WT. We measured P67L interactions using TMT labeled LC-MS proteomics. Overall, we identified 86 statistically significant P67L interactors when comparing the enrichment to the mock control (**Figure 3A**). Similar to F508del, we found upregulated interactors included proteostasis factors such as calnexin (CANX), calmodulin (CALM1), Sec61A, and BAG2 (**Figure 3A**).

**Figure 3.**
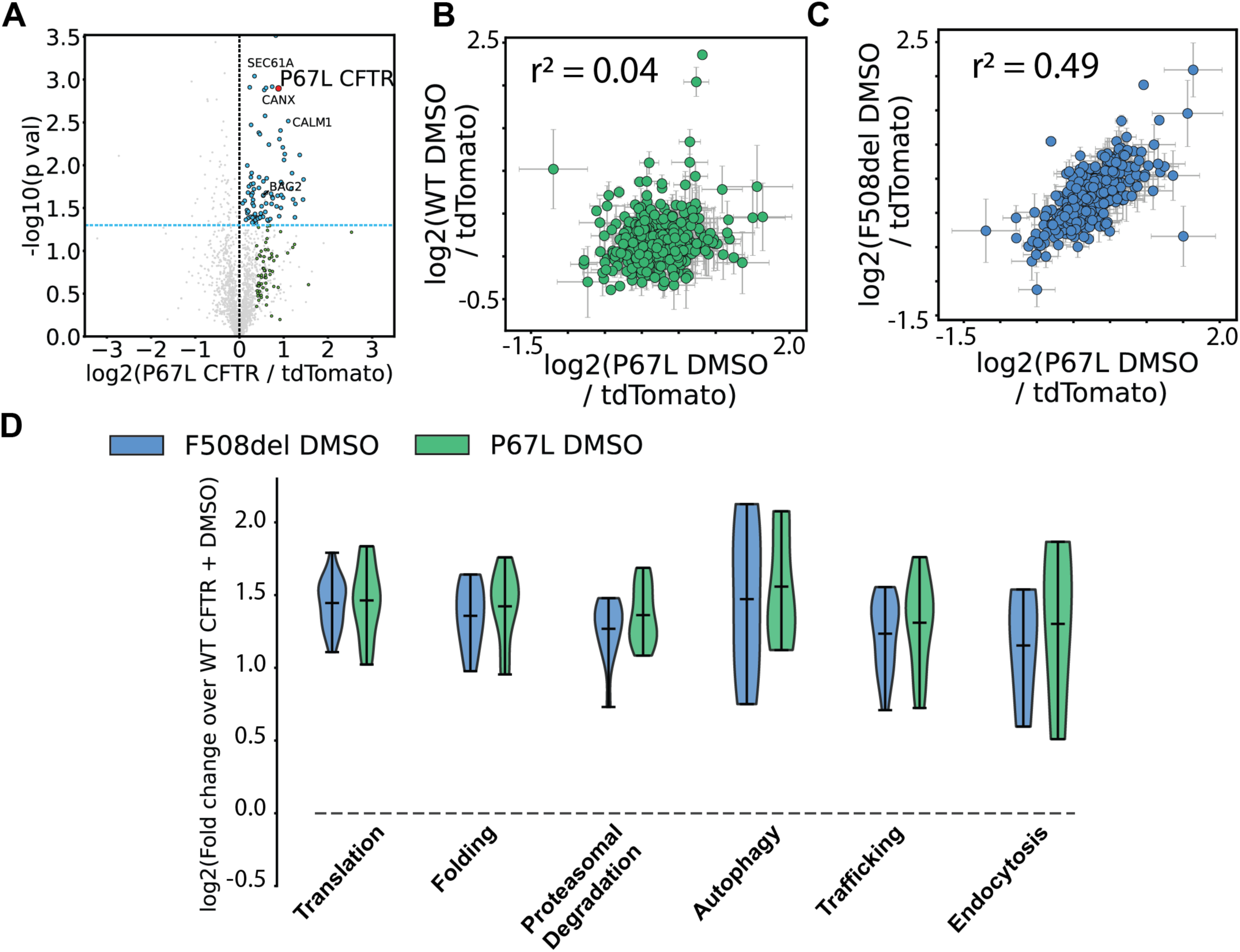
Newly identified P67L interactome correlates with class-II misfolding variant F508del interactome. **A**. Identification of P67L CFTR interactors using multiplexed AP-MS proteomics. The x axis represents log 2-fold change of CFTR IP over IP of tdTomato mock control. A statistical significance cutoff of p > 0.05 is portrayed by a dotted blue line. Blue dots represent newly identified statistically significant interactors in this study and green dots represent previously identified interactors from Pankow, et al.^28^ and Hutt, et al.^35^ that did not meet our statistical cutoff threshold but were enriched greater than one standard deviation of the log 2-fold change distribution between CFTR vs. tdTomato control **B**. Correlation between quantified protein interactors of P67L with DMSO and WT with DMSO. Specific interactions plotted were identified by us or Pankow, et al.^28^ and Hutt, et al.^35^ in our master list. The R-value represents the Pearson correlation coefficient. **C**. Correlation between P67L and DMSO and F508del and DMSO among specific interactions identified by us and others in our master list. **D**. Violin plots comparing the aggregate protein enrichments with important proteostasis pathways for F508del or P67L CFTR. All log2 fold changes are scaled to WT treated with DMSO. Pathways were assigned with GO-terms and annotated manually or by searching previous pathway characterization by Pankow, et al.^28^. These plots demonstrate upregulated interactions with important proteostasis in the class-II variants compared to WT. No statistically significant difference between F508del and P67L was detected for these pathways.

We correlated the interaction fold changes for the class-II variants across all proteins in our master list. Interaction fold changes do not correlate between P67L and WT (Pearson correlation coefficient r^2^ = 0.04) (**Figure 3B**). By contrast, the interaction fold changes correlate well between P67L and F508del (r^2^=0.49) (**Figure 3C**). This is consistent with previous findings that class-II variant interactors correlate better with each other than with WT CFTR^35^. Furthermore, P67L shares more interactors with F508del than with WT (**Supplemental Figure S4A**). Thus, the class-II mutants demonstrate greater overlap with one another than they share with WT. Together these data confirm our hypothesis that the P67L interactome resembles F508del.

Next, we compared the interaction fold changes between F508del and P67L for specific proteostasis pathways with known roles in CFTR processing. For this purpose, we plotted for each pathway the distributions of log2 fold interaction changes for F508del and P67L CFTR scaled to WT (**Figure 3D**). Consistent with the global correlations, both F508del and P67L showed increased interactions with all pathways included in the analysis. P67L demonstrated no substantial differences from F508del in these aggregated interactions. In line with their class-II misfolding categorization, P67L and F508del associated on average more with translational proteins, folding machinery, degradation pathways, and trafficking factors compared to WT (**Figure 3D**). Both mutants also associate more with endocytosis proteins consistent with CFTR quality control at the plasma membrane^51^ (**Figure 3D**). Together, these data demonstrate P67L and F508del share similar increased interactions with important proteostasis pathways.

Overall, our TMT based proteomics workflow successfully captures the proteostasis network engaged by the *hyper-responsive* class-II misfolding CFTR mutation P67L. We found P67L shares specific common interactors with F508del, as well as similar association with proteostasis pathways.

### VX-809 alters P67L interactions at an inflection point between folding and degradation

We next sought to determine how the corrector VX-809 remodels the proteostasis network of P67L CFTR. Specifically, we compared these alterations to F508del to better understand how the proteostasis network influences the *hyper-responsiveness* and trafficking restoration of P67L to the corrector drug. We identified 104 P67L interactors under VX-809 treatment when normalized to tdTomato control (**Figure 4A**). These included previously identified proteostasis factors important for CFTR processing such as BAG2, calnexin (CANX), and calmodulin (CALM1) (**Figure 4A**). Of our identified interactors, 27 of these interactors overlapped with P67L with DMSO and 14 interactors overlapped with WT (**Supplemental Figure S4B**). Thus, P67L treated with VX-809 resembles WT interactome more than P67L with DMSO.

**Figure 4.**
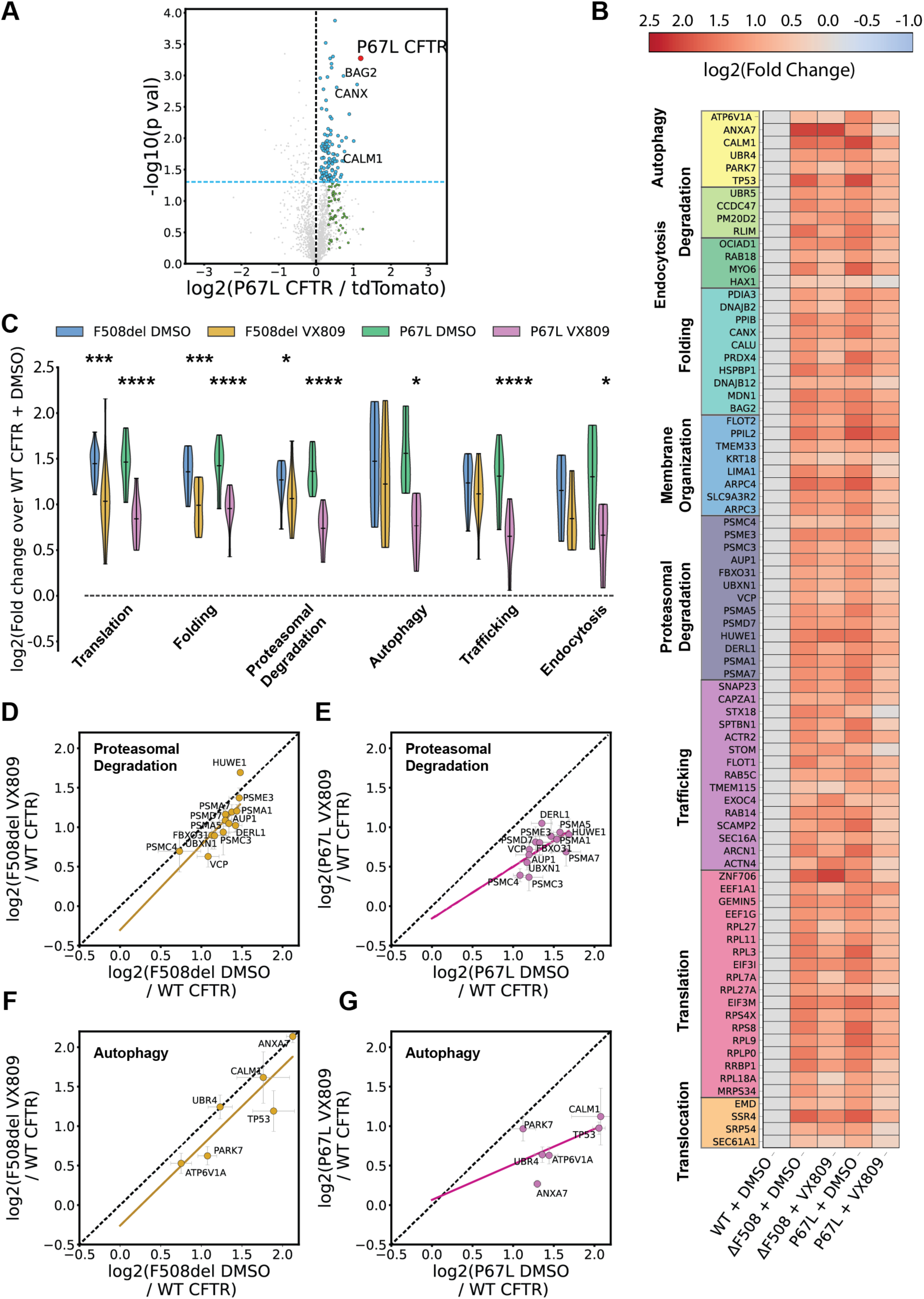
Comparison of *hyper-responsive* P67L and *hypo-responsive* F508del interactomes reveals VX-809 changes pathway interactions at an inflection point between folding and degradation. **A**. Identification of P67L interactors with 3 μM VX-809 treatment using multiplexed AP-MS proteomics. The x axis represents log 2-fold change of CFTR IP over IP of tdTomato mock control. A statistical significance cutoff of p > 0.05 is portrayed by a blue dotted line. Blue dots represent newly identified statistically significant interactors in this study and green dots represent previously identified interactors that did not meet our statistical cutoff threshold but were enriched greater than one standard deviation of the log 2-fold change distribution between CFTR vs. tdTomato control. **B**. A pathway ordered heatmap comparing F508del and P67L interactions under DMSO and VX-809 conditions. Log2 fold changes are scaled to WT with DMSO. Proteins were selected from manually curated list of proteostasis interactors and pathways were assigned with GO-terms. **C**. Violin plots comparing the aggregate protein enrichment levels of important proteostasis pathways for F508del and P67L in the presence and absence of VX-809 treatment (3µM). Pathways were assigned with GO-terms and annotated manually or by searching previous pathway characterization by Pankow, et al.^28^. Violin plots reveal 3 μM VX-809 reduces P67L interactions at an inflection point between proteasomal degradation and autophagy, as these pathways are not as reduced in F508del. Statistical significance calculated by one-way ANOVA with Geisser-Greenhouse correction and Tukey post-hoc multi-hypothesis correction (**Supplemental Dataset 4**). Adjusted p-values depicted by * < 0.05, ** < 0.01, *** < 0.001, and **** < 0.0001. **D**. Proteasomal degradation protein quantifications correlated for F508del between DMSO and 3 μM VX-809 treatment (all log 2 fold changes normalized to WT with DMSO). Black dotted line represents the normal line with a slope of 1 and intercept at the origin. The gold dotted line is the linear least squared best fit for the proteasomal proteins. **E**. Proteasomal degradation protein quantifications correlated for P67L between DMSO and 3 μM VX-809 treatment normalized to WT with DMSO. The purple dotted line is the linear least squared best fit of the proteasomal proteins. **F**. Autophagy protein quantifications correlated for F508del between DMSO and 3 μM VX-809 treatment normalized to WT with DMSO. The gold dotted line is the linear least squared best fit of the autophagy proteins. **G**. Autophagy protein quantifications correlated for P67L between DMSO and 3 μM VX-809 treatment normalized to WT with DMSO. The purple dotted line is the linear least squared best fit of the autophagy proteins.

Next, we quantitatively compared protein levels of P67L interactors between DMSO and VX-809 conditions at the pathway level (**Figure 4B**). The heatmap confirmed the overall reduction in P67L interactors when treated with VX-809. Specifically, VX-809 reduced P67L interaction with calnexin (CANX), BAG2, HSPBP1, DNAJB12, Derlin1 (DERL1), various proteasomal subunits (PSMA1, PSMA5, PSMA7, PSMC3, PSMC4, and PSME3), as well as eukaryotic initiation factors (EIFs) and ribosomal proteins (**Figure 4B**).

To gain insights into how specific proteostasis pathways are impacted by VX-809, we considered the aggregate distribution of interaction changes for individual pathways (**Figure 4C**). We plotted pathway changes in approximate order of biogenesis: translation, folding, proteasomal degradation, autophagy, trafficking, and endocytosis – with the caveat that many of these processes are known to occur simultaneously^52,53^. VX-809 significantly reduced both F508del and P67L interactions with folding and translational machinery with a similar mean change (**Figure 4C**).

Considering later pathways, VX-809 reduced P67L interactions more than F508del interactions with proteasomal degradation and autophagy machinery (**Figure 4C**). Yet, downstream pathways such as trafficking, and endocytosis are only reduced by VX-809 in P67L and not F508del. These data suggest VX-809 reduces P67L proteostasis interactions at an inflection point between folding and proteasomal degradation/autophagy (**Figure 4C**). By contrast, VX-809 does not reduce F508del proteostasis interactions at this inflection point.

Thus, distinct proteostasis alterations emerge when comparing *hypo-responsive* and *hyper-responsive variants*: *the hypo-responsive* F508del continues to experience greater proteasomal and autophagy degradation under VX-809 treatment, while these pathways are attenuated for P67L (**Figure 4C**). Subsequent pathways such as trafficking, and endocytosis also have reduced P67L interactions when treated with VX-809.

To determine which specific proteasomal interactions are reduced by VX-809, we plotted the correlation of interaction fold changes for proteasomal interactors between DMSO vs. VX-809 treatment. We examined these correlations separately for F508del and P67L and fitted each correlation using a linear regression (**Figure 4D-E**). F508del proteasomal interactors remain close to the normal line (black dotted), implying good correlation between DMSO and VX-809 conditions (**Figure 4D**). On the other hand, P67L proteasomal interactors fall below the normal line, implying they are reduced in the VX-809 condition (**Figure 4E**). Interaction with proteasomal subunits PSMC3 and PSMC4 are reduced the most as they fall the furthest beneath the normal line (**Figure 4E**).

Next, to determine which specific autophagy interactions are reduced by VX-809, we plotted the same correlation for these proteins (**Figure 4F-G**). Again, F508del autophagy interactors remain close to the normal line, implying good correlation between DMSO and VX-809 conditions (**Figure 4F**). By contrast, P67L autophagy interactors are globally reduced by VX-809 treatment (**Figure 4G**). In particular, interactions with ANXA7, a GTPase with regulatory role for autophagy^54^, are greatly decreased for P67L in the presence of VX-809 (**Figure 4G**). These analyses highlight other potential protein targets for CF treatment in less responsive mutants. We also compared correlations for translation and folding proteins (**Supplemental Figure S4C-D**). These plots reveal further interactions of P67L that are greatly attenuated with VX-809 treatment, such as DNAJB12 in the folding correlations (**Supplemental Figure S4D**). Together, these data highlight that VX-809 alters specific proteostasis interactions for P67L compared to F508del that may underlie the *hyper-response* to the corrector.

In summary, our data shows commonalities in the way that VX-809 treatment attenuates the proteostasis interactions of both F508del and P67L, reducing interactions with folding and translation proteins. At the same time, distinct interaction changes between the two mutants point to routes by which VX-809 treatment can better restore P67L plasma membrane trafficking. Our data indicate that resetting of degradation interactions may represent an inflection point in the *hyper-responsiveness* for P67L as these pathways are reset closer to WT-levels for P67L than for F508del (**Figure 4C**). Subsequently, P67L also experiences a reduced interaction with trafficking machinery under VX-809 treatment, which could indicate faster flux through the pathway.

### L206W and G85E CFTR interactomics reveals pathway commonalities between hyper-responsive and hypo-responsive variants

To further understand how VX-809 treatment generally affects CFTR interactions, we turned to another *hyper-responsive* mutation, L206W, as well as the *hypo-responsive* mutant G85E. L206W is an N-terminal class-II CFTR variant associated with mild CF disease^55^ (**Figure 1A**). L206W impairs CFTR biosynthesis, typical of class-II variants, but demonstrates single-channel conductance similar to WT CFTR^56^ – indicating that plasma membrane-localized L206W retains some function. Second, we chose G85E, an N-terminal class-II variant that cannot be corrected by VX-809^1^ (**Figure 1A**).

L206W CFTR exhibited near WT-levels of band C when treated with VX-809, confirming a *hyper-response* like P67L (**Figure 5A**). On the other hand, very minimal band C CFTR was observed for the G85E variant regardless of VX-809 addition (**Figure 5B**). We measured L206W and G85E CFTR protein interactions with TMT labeled LC-MS/MS to understand what common and distinct interaction changes occur with VX-809 treatment for the *hyper-* and *hypo-responsive* variants. Importantly, several of the TMT11plex sets included conditions for WT CFTR and all four variants (+/- VX-809) allowing for direct comparison of the interaction fold changes in the integrated dataset (**Supplemental Dataset 1**).

**Figure 5.**
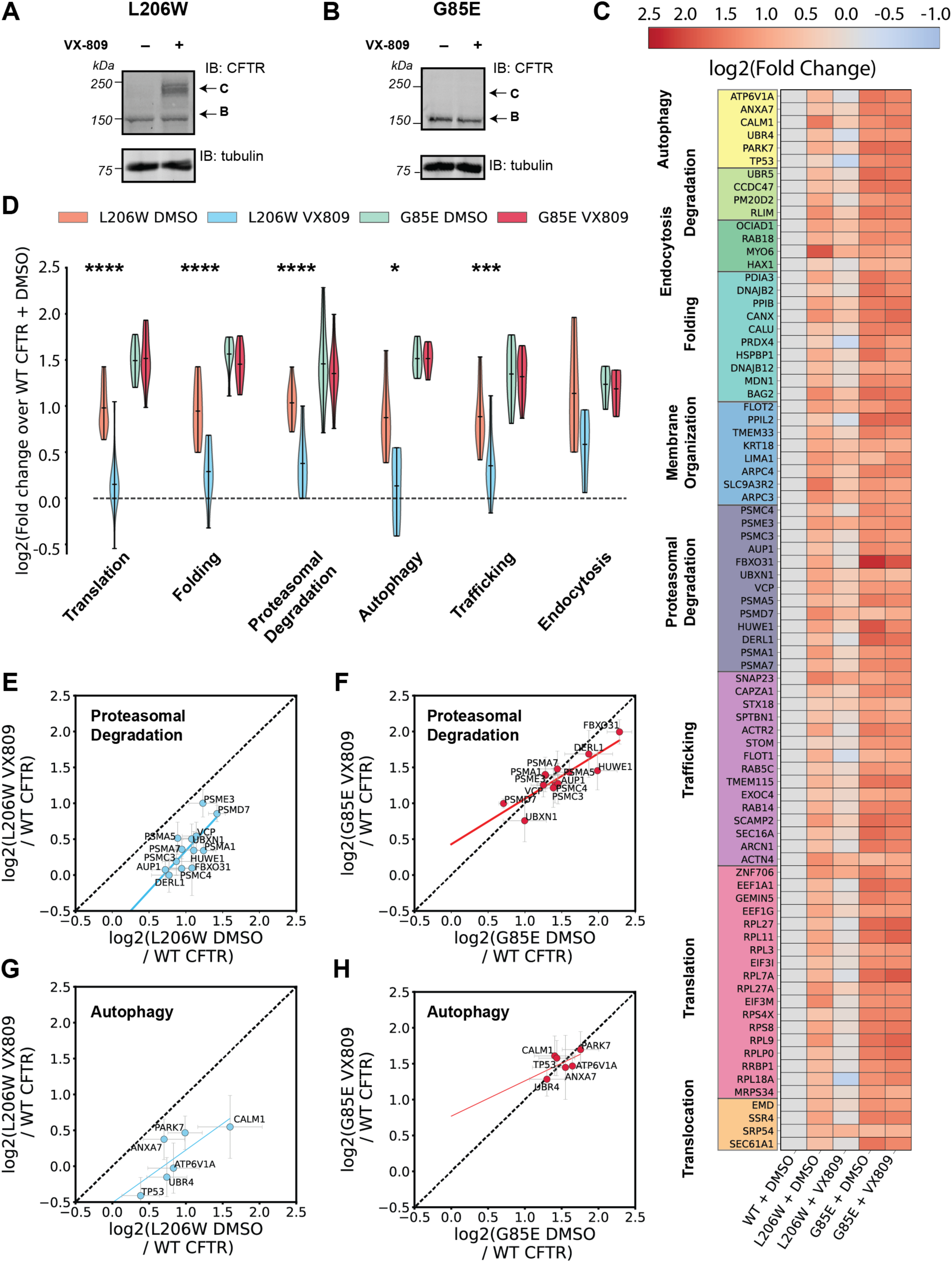
L206W interactome with VX-809 reveals similar changes in degradation pathways not observable in non-responsive G85E. **A-B**. Western blot cell lysates expressing L206W (**A**) or G85E (**B**) CFTR probed with CFTR 217 antibody and tubulin loading control. Comparison of C Band CFTR between DMSO and 3 μM VX-809 treatment demonstrates L206W classification as a *hyper-responsive* variant, while G85E is a *non-responsive* variant. **C**. A pathway ordered heatmap comparing L206W and G85E interactions under DMSO and 3 μM VX-809 conditions (all log2 fold changes scaled to WT with DMSO). Proteins were selected from manually curated list of proteostasis interactors and pathways were assigned with GO-terms. **D**. Violin plots comparing aggregate protein enrichment levels for important proteostasis pathways between L206W and G85E CFTR in the presence and absence of VX-809 (3 µM). Pathways were assigned with GO-terms and annotated manually or by searching previous pathway characterization by Pankow, et al.^28^. Violin plots reveal 3 μM VX-809 reduces L206W proteostasis interactions which are not reduced in G85E. (**Supplemental Dataset 4**). Statistical significance calculated by one-way ANOVA with Geisser-Greenhouse correction and Tukey post-hoc multi-hypothesis correction. Adjusted p values depicted by * < 0.05, ** < 0.01, *** < 0.001, and **** < 0.0001. **E**. Proteasomal degradation protein quantifications correlated for L206W between DMSO and 3 μM VX-809 treatment normalized to WT with DMSO. Black dotted line represents a normal line with a slope of 1 and intercept at the origin. The blue dotted line is the linear least squared best fit of the proteasomal proteins. **F**. Proteasomal degradation protein quantifications correlated for G85E between DMSO and 3 μM VX-809 treatment normalized to WT with DMSO. The red dotted line is the linear least squared best fit of the proteasomal proteins. **G**. Autophagy protein quantifications correlated for L206W between DMSO and 3 μM VX-809 treatment normalized to WT with DMSO. The blue dotted line is the linear least squared best fit of the autophagy proteins. **H**. Autophagy protein quantifications correlated for G85E between DMSO and 3 μM VX-809 treatment normalized to WT with DMSO. The red dotted line is the linear least squared best fit of the autophagy proteins.

We first defined statistically significant interactors of the individual L206W and G85E variants. A previous proteomics dataset has been published on G85E for comparison^35^. We identified 86 statistically significant L206W interactors under DMSO and 66 interactors under VX-809 treatment (**Supplemental Figure S5A-S5B**). We also identified 206 statistically significant G85E interactors under DMSO and 146 interactors under VX-809 treatment (**Supplemental Figure S5C-S5D**), consistent with an increased number of interactions with *hypo-responsive* variants seen in a previous published study^28^. From our dataset, 27 G85E interactors overlapped with the previous G85E datasets, albeit the general overlap was relatively small (13 %) (**Supplemental Figure S5E**).

Next, we compared the interaction fold changes between CFTR variants under VX-809 treatment conditions (**Figure 5C**). As with the prior class II CFTR mutant, we observed overall increases in proteostasis interactions with L206W and G85E compared to WT when treated with DMSO. This is consistent with the enhanced proteostasis surveillance experienced by these destabilized, trafficking-defective mutants. The overall magnitude of the increase is lower for L206W compared for G85E (**Figure 5C**), which could reflect the milder trafficking defect of L206W. We then examined how VX-809 treatment restores the proteostasis interactions. The data clearly highlighted that interactions decrease globally for L206W under VX-809 treatment. By contrast, G85E demonstrated little change in the presence of the corrector.

To gain deeper insight into the impact of VX-809 treatment on distinct proteostasis pathways, we plotted the interaction abundance changes grouped by pathways under the various conditions (**Figure 5D**). Like P67L, L206W experienced significantly decreased interactions with translational, folding, proteasomal degradation, autophagy, and trafficking pathways. On the other hand, G85E did not show a significant change in any pathway interactions in the presence of VX-809, which reflects its classification as a *hypo-responsive* variant (**Figure 5D**). Together, these data highlight that pathway analysis of interaction changes for CFTR variants under VX-809 treatment can clearly distinguish *hyper-responsive* and *hypo-responsive* CFTR variants.

To elucidate specific proteasomal interactions reduced by VX-809, we plotted the correlation of interaction fold changes for proteasomal interactors with DMSO vs. VX-809 treatment for L206W. We compared this correlation to G85E (**Figure 5E-F**). Linear regression best fit reveals L206W proteasomal interactions fall below the normal line, implying VX-809 reduces these interactions (**Figure 5E**). L206W interactions with proteasomal proteins are globally reduced and reveal few outliers (**Figure 5E**). By contrast, G85E proteasomal interactions straddle the normal line, implying VX-809 does not affect these interactions (**Figure 5F**). Likewise, L206W autophagy interactions fall below the normal implying reduction by VX-809 (**Figure 5G**), whereas G85E interactions remain unaffected (**Figure 5H**). Furthermore, L206W interactions fall below the normal for translational, folding, and trafficking proteins, where G85E interactions always fall near the normal line (**Supplemental Figure S6**).

Overall, the data shows that *hyper-responsive* variants, such as P67L and L206W, display global attenuation of proteostasis interactions closer to WT levels, in particular reduced interactions with proteasomal degradation and autophagy pathways. In contrast, *hypo-responsive* variants maintain prolonged interactions with translation machinery and are not rescued from engagement with degradation machinery by VX-809 treatment. An interesting distinction between the two *hypo-responsive* variants is that F508del displays some reductions in proteostasis interactions with folding and translation components when treated with VX-809 (**Figure 4D**), while G85E interactions are completely refractory to the corrector compound (**Figure 5E**). This difference in interaction changes likely reflects the distinct theratype, as G85E is entirely non-responsive to VX-809 treatment while F508del exhibits mild restoration of plasma membrane trafficking and function as observed via Western blot and Ussing chamber studies^3^.

Measuring the interactions of distinct drug responsive variants reveals the defining characteristics of *hyper-responsive* mutations, or theratype. The ability of VX-809 to attenuate interactions with translational, folding, and proteasomal degradation components in responsive CFTR variants appears to be critical to this pharmacological chaperone theratype.

### Inhibiting proteasomal degradation and autophagy differentially impacts the VX-809 rescue of hyper-responders and hypo-responders

We demonstrated VX-809 likely rescues *hyper-responsive* CFTR variants P67L and L206W by reducing interactions at an inflection point in biogenesis involving proteasomal degradation and autophagy machinery. On the contrary, interactions with these degradation pathways were not as reduced for the *hypo-responsive* variants F508del and G85E. For this reason, we chose to further investigate how proteasomal and autophagy degradation pathways contribute to the divergent VX-809 correction levels experienced by the *hyper-* and *hypo-responsive* CFTR variants. F508del is a well-known target of ER-associated degradation (ERAD) which directs the protein to the proteosome^57^. Inhibition of ERAD by proteosome inhibitors such as bortezomib^58^ is known to partially rescue F508del CFTR^46^.

To test the impact of proteasome inhibition on VX-809 rescue, cells expressing mutant CFTR variants were co-treated with VX-809 and bortezomib. Subsequent effects on CFTR expression and trafficking were analyzed by Western blot. Bortezomib treatment alone resulted in a strong 4-fold increase in F508del band A (immature, non-glycosylated) and band B (core-glycosylated, ER localized) CFTR (**Figure 6A-B**). Bortezomib also led to a 3-fold increase in mature band C CFTR, and this increase was slightly higher than with VX-809 treatment alone (**Figure 6A, 6C**). These data demonstrated that proteasome inhibition alone can restore F508del maturation, which is consistent with prior studies^46^. We also quantified trafficking efficiency by considering the ratio of band C band relative to total CFTR (band C/[band B + band C]) (**Supplemental Figure S7A**). Trafficking efficiency remains similar for F508del when cells were treated with bortezomib or DMSO, highlighting that some of the CFTR buildup in the ER is trafficking competent and subsequently increases a similar proportion of mature CFTR in the plasma membrane. We then compared how bortezomib treatment affects the VX-809 response for F508del. Co-treatment resulted in higher band C F508del amounts compared to bortezomib or VX-809 treatment alone (**Figure 6A, C**). Trafficking efficiency continued to be increased compared to the untreated condition, albeit slightly lower than when treated with VX-809 alone, indicating that proteasome inhibition does not completely impair VX-809 correction but still attenuates it (**Supplemental Figure S7A**).

**Figure 6.**
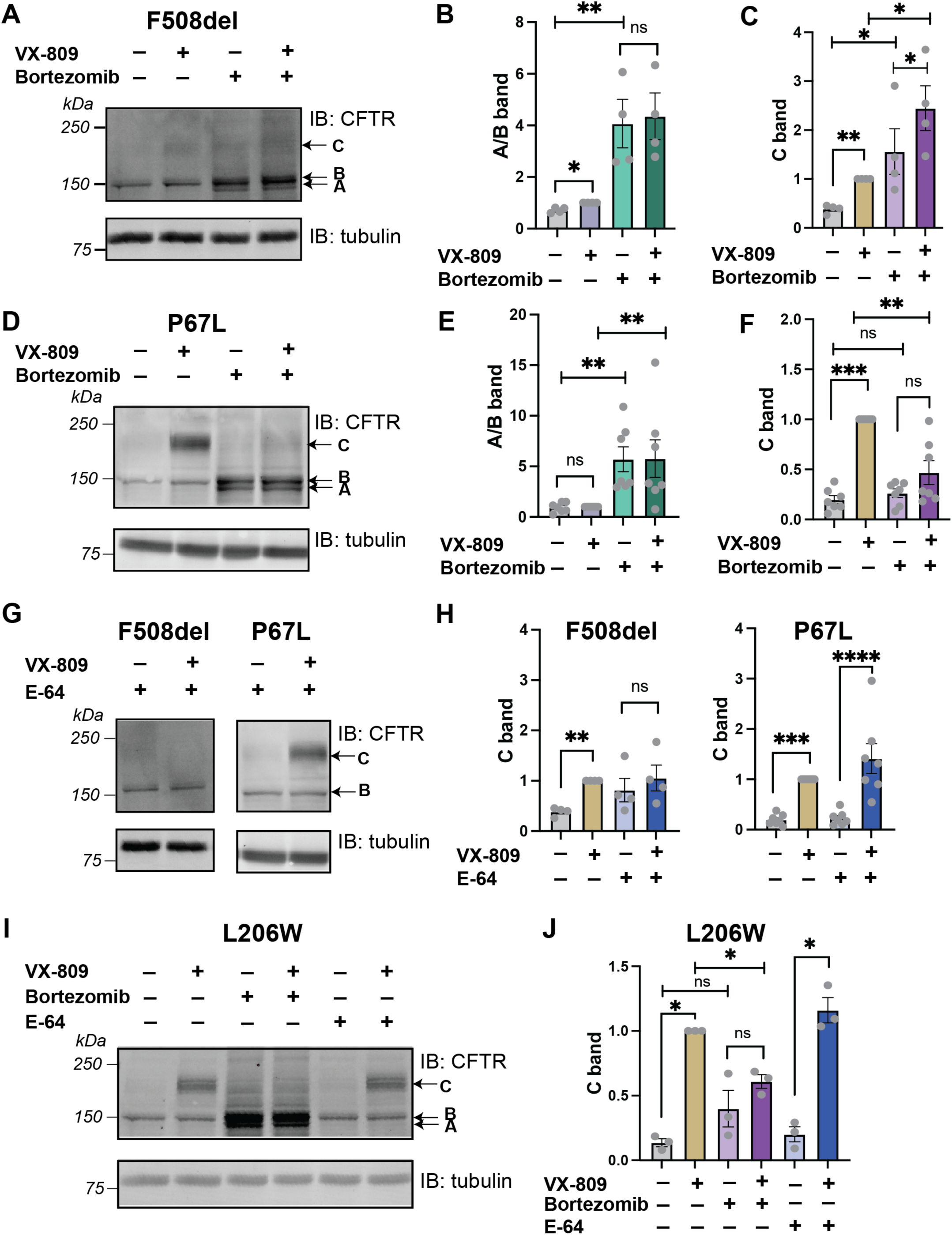
Inhibiting proteasomal degradation attenuates *hyper-responsive* P67L and L206W VX-809 response, while inhibiting lysosomal degradation attenuates *hypo-responsive* F508del VX-809 response. **A**. Representative Western blot of F508del CFTR trafficking assay. Cells expressing F508del CFTR were treated with 3 μM VX-809, 10 μM bortezomib, or a combination. Cell lysates samples were separated by SDS-PAGE and blots were probed with 217 CFTR antibody and tubulin antibody as a loading control. **B**. Quantification of immature (A/B band) F508del levels from western blot data. Protein bands for band A/B F508del CFTR were normalized to the 3 μM VX-809 treatment condition. **C**. Quantification of mature (C band) F508del levels from western blot data showing. Band C (post-Golgi) F508del CFTR levels were normalized to the 3 μM VX-809 treatment condition. **D**. Representative Western blot of P67L CFTR trafficking assay. Cells expressing F508del CFTR were treated with 3 μM VX-809, 10 μM bortezomib, or a combination. Blots were probed with 217 CFTR antibody and tubulin antibody as a loading control. **E**. Quantification of immature (A/B band) P67L western blot data showing the relative ER retained P67L CFTR normalized to the 3 μM VX-809 treatment condition. **F**. Quantification of mature (C band) P67L western blot data showing the relative post-Golgi P67L CFTR normalized to the 3 μM VX-809 treatment condition. **G**. Representative Western blot of F508del and P67L CFTR trafficking assay when cells were treated 10 μM E-64 (cysteine protease inhibitors) or a combination of E-64 with 3 μM VX-809. Blots were probed with 217 CFTR antibody and tubulin antibody as a loading control. **H**. Quantification of mature (C band) CFTR western blot data for F508del and P67L showing the relative post-Golgi CFTR normalized to the 3 μM VX-809 treatment condition. **I**. Representative Western blot of L206W CFTR trafficking assay when cells were treated with 3 μM VX-809, 10 μM bortezomib, 10 μM E-64, or a combination. Blots were probed with 217 CFTR antibody and tubulin antibody as a loading control. **J**. Quantification of mature (C band) L206W CFTR western blot data showing the relative post-Golgi CFTR normalized to the 3 μM VX-809 treatment condition. For all graphs, individual measurements are shown as grey points, error bars represent standard error of the mean, statistical significance was calculated with a paired, two tailed student t-test and p-values are depicted by * < 0.05, ** < 0.01, *** < 0.001 and **** < 0.0001.

We then probed the impact of proteasome inhibition on P67L CFTR. Bortezomib treatment resulted in more than 5-fold greater band B P67L accumulation (**Figure 6D-E**). However, in contrast to F508del, proteasome inhibition did not increase band C P67L CFTR (**Figure 6D, F**), suggesting that the accumulated immature P67L population does not traffic. We then tested whether VX-809 could restore P67L trafficking in the presence of proteasome inhibition. Surprisingly, co-treatment of bortezomib and VX-809 abrogated P67L band C accumulation (**Figure 6D, F**), indicating that the proteasome inhibitor interferes with VX-809 correction. Similarly, co-treatment of bortezomib and VX-809 reduced trafficking efficiency as measure by the band C/(band B + band C) ratio compared to VX-809 correction alone (**Supplemental Figure S7B**). These data further suggest a previously unknown role for proteasomal degradation in VX-809 corrections.

Given the distinct impact of proteasome inhibition on P67L and F508del, we turned to examine additional degradation pathways. Auto-lysosomal degradation is another major pathway for removal of CFTR from cells. Multiple routes deliver CFTR to the autolysosome, including ER-associated autophagy and endocytic pathways removing CFTR from the plasma membrane^51,59^. We used the cathepsin protease inhibitor E-64 to broadly block lysosomal protein degradation^60^. E-64 treatment did not result in any accumulation of F508del or P67L band B or band C (**Figure 6G-H, Supplemental Figure S7C-D**), confirming that auto-lysosomal degradation does not represent a major pathway in the clearance of these variants. We then co-treated E-64 with VX-809 to probe its impact on the VX-809 correction mechanism. For P67L, the lysosomal protease inhibitor had no effect on VX-809 correction as seen by similarly increased band C levels (**Figure 6G-H**) and increased trafficking efficiency (**Supplemental Figure S7B**) compared to VX-809 treatment alone. In contrast, E-64 abrogated the small degree of VX-809 correction for F508del (**Figure G-H, Supplemental Figure S7A**). These results indicate that autolysosomal degradation is necessary for the VX-809 correction of F508del, but not for P67L.

The divergent dependencies of the two CFTR variants on degradation pathways during VX-809 correction prompted us to turn to the L206W and G85E variants. Bortezomib treatment resulted in buildup of band B CFTR for all variants (**Figure 6I, Supplemental Figure S7E, G, H**). On the other hand, E-64 addition did not lead to any accumulation of band B or C CFTR (**Figure 6I-J, Supplemental Figure S7G-I**), demonstrating that degradation of those variants also primarily occurs through ERAD. Co-treatment of inhibitors with VX-809 showed that proteasomal inhibition again impaired the correction of L206W, but auto-lysosomal inhibition did not (**Figure 6I-J, Supplemental Figure S7F)**. Thus, both *hyper-responsive* L206W and P67L variants exhibit similar dependences on active proteasomal degradation for correction by VX-809. Lastly, G85E exhibited a small amount of restoration of band C CFTR with bortezomib treatment regardless of VX-809 treatment (**Supplemental Figure S7G, I**). Like F508del, this increased expression likely stems from increased ER retained G85E CFTR as the trafficking efficiency of G85E is unaffected (**Supplemental Figure S7J**). However, VX-809 did not lead to any further correction, confirming that G85E is completely refractory to the corrector compound. Together these data suggest that auto-lysosomal inhibition does not affect VX-809 *hyper-responders* P67L and L206W but impairs F508del VX-809 response. In contrast, we determined that proteasomal degradation is a necessary pathway for the effective correction of the *hyper-responsive* variants.

## DISCUSSION

This study represents, to our knowledge, the first quantitative interactomics analyses comparing CFTR mutations with diverse VX-809 response. We used AP-MS coupled to TMT labeling, which allows sample multiplexing and direct quantitative comparison of protein interaction changes between mutants and treatments simultaneously. Prior studies from our group used this platform to explore how the endoplasmic reticulum proteostasis network (ER PN) facilitates protein quality control for misfolding-prone proteins, such as amyloidogenic light chains and thyroid prohormones^40,61^. Here, we extend the multiplexed, quantitative TMT-based mass spectrometry platform to compare CFTR interactome changes across patient mutations F508del, P67L, L206W, and G85E, which exhibit distinct response profiles to clinically approved therapeutic correction. Comparing drug-induced interaction changes between *hyper-responsive* mutations (P67L and L206W) and *hypo-responsive* mutations (F508del and G85E) revealed important proteostasis components that may drive drug susceptibility. Specifically, we found *hyper-responsive* mutations to associate less with proteasomal and autophagy degradation machinery.

We first evaluated the ability of our approach to recapitulate the known proteostasis interactions of F508del CFTR. Consistent with previous findings, we show F508del interacts more with proteostasis factors than WT. We identified many novel F508del interactors as well as demonstrated comparable overlap with previous data^28,35^. Importantly, our TMT-based quantification provides the ability to directly compare F508del interactions under VX-809 treatment. This analysis revealed that F508del interacts less with proteostasis components when treated with VX-809 similar to F508del interactome remodeling during temperature correction^28^. Specifically, VX-809 reduced F508del interactions with translational, folding, and proteasomal degradation pathways. On the individual protein level, VX-809 lowered interactions with previous characterized F508del proteostasis components such as calnexin (CANX), MYO6, BAG2, DNAJB12, DNAJB2 and several factors whose knockdown restores F508del trafficking. These data suggest our TMT quantification approach can measure differences in F508del interactions under drug treatment with proven roles in CFTR protein quality control. Thus, we turned to P67L to determine what protein interactions underlie *hyper-responsive* VX-809 correction.

Despite extensive characterization of P67L as a VX-809 *hyper-responsive* CFTR variant^19,50^, its proteostasis interactome remained undefined until this study. We found increased overlap and correlation with F508del compared to WT CFTR. These overlapping proteostasis components included chaperones, degradation machinery, translation, and trafficking proteins. We then quantitatively compared P67L interactions under VX-809 treatment conditions to determine which specific protein interaction changes are associated with the *hyper-responsive* behavior of P67L CFTR to the corrector drug. In the presence of VX-809, the P67L interactions resembled WT levels. This was evident by greater overlap in interactions with WT CFTR than with P67L + DMSO, and a broad reduction in the magnitude of interactions with proteostasis factors. These observations are consistent with a shift towards WT protein quality control for *hyper-responsive* mutations during correction. Commonalities for P67L and F508del CFTR are attenuated interactions with translation and folding components in the presence of VX-809. However, an important finding was that VX-809 uniquely attenuates interactions with proteasomal degradation, autophagy, trafficking, and endocytosis proteins for P67L. These interactions are maintained for F508del even in the presence of VX-809, pointing towards their distinct role in the *hyper-responsive* behavior. To determine the interaction commonalities underlying robust VX-809 response, we followed up by determining the,interactome changes of two other mutants: *hyper-responsive* L206W – which represents a novel interactome not previously published, and *hypo-responder* G85E. We showed VX-809 reduced L206W interactions globally across many pathways when compared to WT CFTR – confirming VX-809 correction remodels a near-WT interactome. By contrast, VX-809 does not appear to alter the G85E interactome in any pathway.

What can *hyper-responsive* CFTR mutations reveal about the VX-809 mechanism? We showed P67L and L206W experience reduced interaction with degradation machinery and pathways downstream of folding compared to F508del. VX-809 corrects both full length F508del CFTR at the plasma membrane^62^ and nascent F508del CFTR during biogenesis^3^. Our data suggests L206W and P67L are likely corrected by VX-809 binding early in biogenesis – consistent with proposed binding sites at TMD1^63^ and the TMD1/NBD1 interface^64^. Importantly, since CFTR degradation and folding occur co-translationally^52,53^ and likely before trafficking, we speculate reduction in trafficking proteins occur after reduction of degradation machinery. Thus, we propose a model where VX-809 primarily acts at an inflection point in biogenesis before routing towards proteasomal degradation (**Figure 7A**).

**Figure 7.**
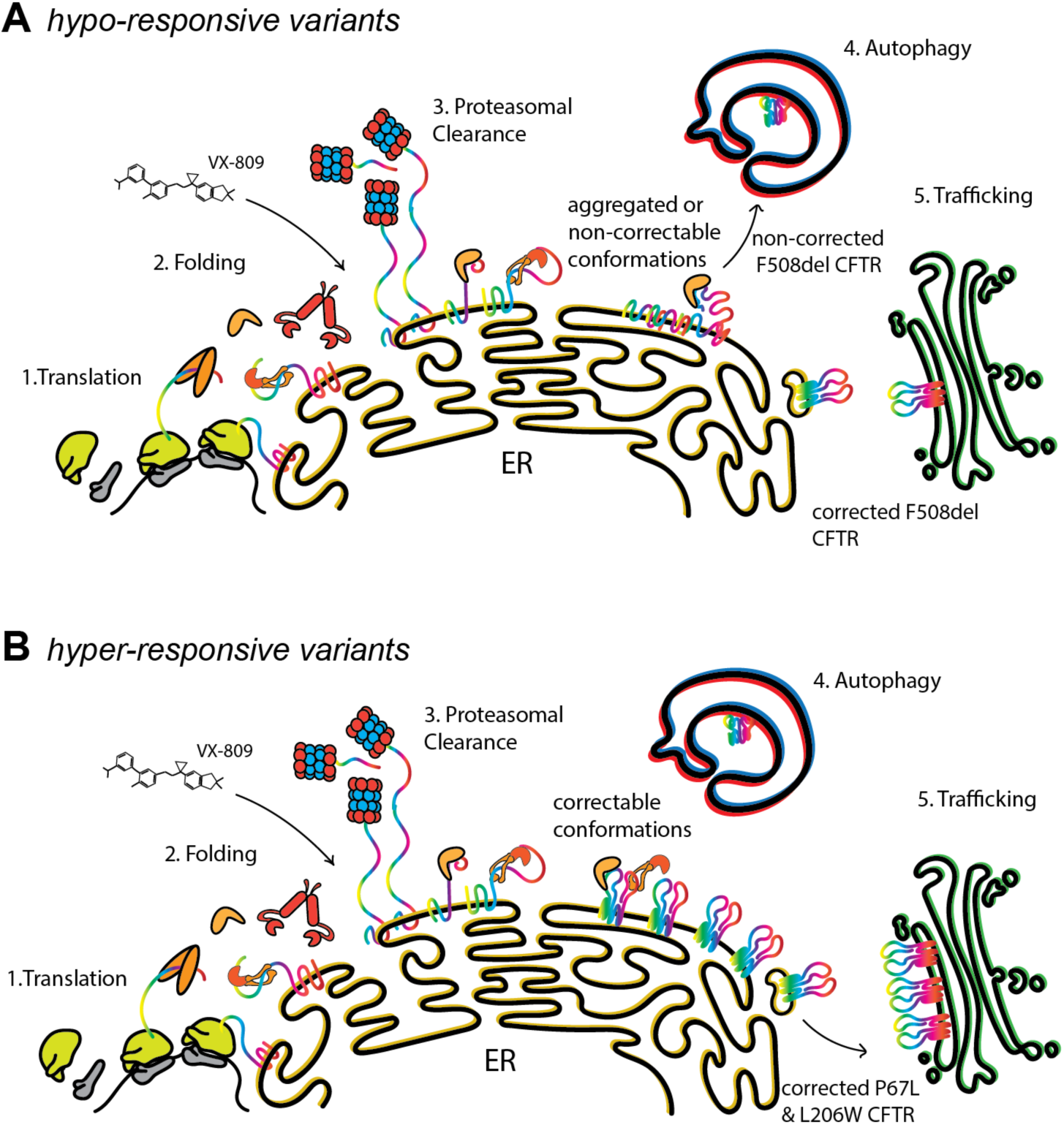
Model of VX-809 acting an inflection point just prior to protein degradation that requires clearance of non-correctable CFTR conformations. **A**. Schematic indicating the protein quality control process for *hypo-responsive* variant F508del. VX-809 reduces F508del interactions with translation and folding proteins, however, not interactions with proteasomal degradation and autophagy proteins, and thus is likely acting at an inflection point between folding and proteasomal degradation. Since inhibiting autophagy attenuates F508del VX-809 response, it is likely that autophagy clearance of non-correctable conformations is required for VX-809 to correct F508del. **B**. Schematic showing fates of *hyper-responsive* variants P67L and L206W. VX-809 reduced P67L and L206W interactions with translation, folding, proteasomal degradation, autophagy and trafficking proteins allowing robust VX-809 response indicative in the increased trafficking. Since inhibiting proteasomal degradation attenuates P67L and L206W VX-809 response, it is likely that proteasomal clearance of non-correctable conformations is required for VX-809 to correct P67L and L206W.

Recently, He, et al. showed N1303K CFTR is degraded by ER-associated autophagy involving DNAJB12 and they speculated N1303K may not be retro-tranlocatable^59^. By contrast, they showed P67L/N1303K CFTR is not detected in autolysosomes and like degraded proteasomally^59^. However, when treated with VX-809, P67L/N1303K is reverted back to ER-associated autophagy similar to N1303K^59^. This suggests P67L is corrected by VX-809 at a step before auto-lysosomal degradation pathways (**Figure 7B**). However, precisely when this step occurs during CFTR folding remains unclear. We previously showed the P67L VX-809 response requires the CFTR C terminus^50^. Chaperones such as DNAJB12 evaluate CFTR N-terminal/C-terminal interactions.^65^ DNAJB12 has been shown to triage F508del CFTR through proteasomal degradation^66,67^ and N1303K CFTR through autophagy^59^. Indeed, we show that P67L-DNAJB12 interactions are particularly reduced among folding factors by VX-809 (**Supplemental Figure 4B**). Furthermore, Hsp40 and Hsp70 chaperones bind the TMD2/lasso motif interface^27^ near P67L. Thus, we speculate VX-809 rescues P67L during a chaperone engaged conformation that is still susceptible to retro-translocation and proteasomal degradation. This likely occurs when N-terminal/C-terminal interactions between the lasso motif/TMD2 are forming – which are perturbed in P67L, but rescued by VX-809^50^.

This is in contrast to F508del CFTR which perturbs both NBD1/TMD2 interactions^68,69^ and disrupts NBD2 post-translational folding^53^. Thus, auto-lysosomal degradation may triage F508del CFTR due to NBD2 folding defects, like N1303K, regardless of VX-809 correction (**Figure 7A**). However, we demonstrated that inhibition of lysosomal protease with E-64 attenuates F508del VX-809 response, therefore the role of lysosomal degradation remains unclear. Thus, we can only speculate on the precise folding events at which VX-809 becomes crucial for *hyper-responsive* mutations, but our data suggests this occurs at an inflection point during or just prior to proteasomal degradation.

We revealed in this study that VX-809 becomes crucial for *hyper-responsive* mutations before proteasomal degradation. Furthermore, we demonstrated that proteasome inhibition attenuates P67L and L206W VX-809 response. These data suggest that proteasomal clearance is required to maintain a steady-state population of VX-809 correctable CFTR (**Figure 7B**). Two alternative models may explain this observation. First, mutant CFTR may be more prone to aggregation, thus requiring proteasomal clearance for a steady state correctable population. This correctable CFTR population may be chaperone bound, undergoing post-translational folding, and near a critical step in N-terminal/C-terminal interdomain assembly. Alternatively, under proteasome inhibition cells undergo a starvation response and upregulate autophagy^70^. In this model, CFTR conformations bound for autophagy may be correctable and the increase in autophagy would reduce this correctable population and prevent VX-809 response.

VX-809 *hyper-responsive* CFTR variants P67L and L206W share several commonalities. We measured the novel L206W interactome - revealing L206W shares similar proteostasis interactions as other class-II mutations- and the interactome of hypo-responder G85E. We showed VX-809 reduced L206W interactions globally across many pathways indicating VX-809 correction remodels an almost near-WT interactome. By contrast, VX-809 does not appear to alter the G85E interactome in any pathway. On a pathway specific basis, VX-809 reduced L206W with folding, translational, proteasomal, autophagy and trafficking machinery. Thus, L206W is likely corrected by VX-809 binding early in biogenesis – consistent with notions of TMD1^63^ and the TMD1/NBD1 interface^64^ binding sites. By contrast, since G85 only interacts with residues in TMD1 (**Supplemental Figure S5F**) it is likely that a mutation here experiences misfolding early in biogenesis and cannot reach a correctable conformation by slowing membrane insertion^71^. This is consistent with our Western blots showing that proteasomal inhibition increases G85E C band but does not synergize with VX-809. This suggests that some structural defects may simply be un-correctable, and possibly such defects precede protein interaction alterations.

Thus, VX-809 *hyper-responsive* CFTR mutants most likely reach a correctable conformation before the checkpoint to route for proteasomal degradation. Since VX-809 binding sites remains elusive, mechanistic studies of how the corrector changes CFTR interactions provides valuable information of a possible drug mechanism. Our TMT labeling approach provides a foundation for future work using a variety of CFTR mutations and corrector to theratype mutants.

Besides VX-809, additional pharmacologic corrector drugs are now in clinical use, including the combination treatment Trikafta, which was FDA-approved in 2019. Trikafta contains the potentiator VX-770 (Ivacaftor) and two corrector compounds: VX-661 (Tezacaftor), which is a structural analog of VX-809, and VX-445 (Elexacaftor). The combination treatment is currently approved for patients carrying F508del hetero- and homozygous CFTR variants (nearly 90% of CF patients), however, many rare CFTR variants are not included in this list^5^. Elucidation of the corrector theratypes for rare CFTR mutants and a better understanding of the principles that dictate their susceptibility to individual corrector compounds could broaden their therapeutic use for personalized CF medicine. Our multiplexed approach could facilitate the high-throughput characterization of altered proteostasis interactions of theratypes for future CFTR correctors. Beyond theratyping CFTR mutants, our high-throughput multiplexed TMT quantification approach offers the capability to theratype mutant protein variants associated with other protein misfolding diseases and respective pharmacologic chaperones.

## MATERIALS AND METHODS

### Plasmids and Antibodies

Plasmids expressing CFTR in the pcDNA5 vector were gifted from E. Sorscher and J. Hong (Emory University, Atlanta, Georgia, USA)^19^. Anti-CFTR mouse monoclonal antibodies included 217 R-domain antibodies (http://cftrantibodies.web.unc.edu/) (provided by J. Riordan, University of North Carolina, Chapel Hill, North Carolina, USA) and 24-1 anti–C-terminus monoclonal antibodies (mAB 24-1 (ATCC® HB-11947™) purified from B lymphocyte hybridoma cells using a Recombinant Protein G - Sepharose 4B Affinity Column on an ÄKTA start protein purification system (GE Life Science Product # 29022094). Antibodies were used in immunoblotting buffer (5% bovine serum albumin (BSA) in Tris-buffered saline pH 7.5, 0.1% Tween-20, and 0.1% NaN_3_) with 217 at 1:1000 dilutions. Secondary antibodies were obtained from commercial sources and used at the indicated dilutions in 5% milk in tris-buffered saline, 0.1% Tween-20 (TBS-T): Goat anti-mouse Starbright700 (1:10000, Bio-Rad), anti-rabbit rhodamine-conjugated tubulin (1:10000, Bio-Rad).

### Immunoblotting, SDS-PAGE, and Immunoprecipitation

HEK293T cells were transiently transfected with respective CFTR expression plasmids using a calcium phosphate method^72^. A fully confluent 10cm plate (approximately 10^7^ cells) was used per condition. Cells were harvested by washing with PBS and incubated with 600 µL of TN buffer (50 mM Tris pH 7.5, 150 mM NaCL) with 0.5% IGEPAL CA-630 and complete, Mini, EDTA-free Protease Inhibitor Cocktail (Roche) in PBS and rocked at 4 °C for 20 minutes. A cell scraper was then used to dislodge cells and transfer them to a micro-centrifuge tube. Cell were sonicated for 3 minutes and spun down at 18,000 g for 30 minutes. Total protein concentration was normalized using Bio-Rad protein assay dye. A sample of the lysates (used as the “input”) was then denatured with 6X Laemmli buffer + 100mM DTT and heated at 37°C for 15 minutes before being separated by SDS-PAGE. Samples were transferred onto PVDF membranes (Millipore) for immunoblotting and dehydrated. Primary antibodies were incubated either at room temperature for 2 hours, or overnight at 4°C. Membranes were then washed three times with TBS-T and incubated with secondary antibody constituted in 5% milk at 4°C for 1 hour. Membranes were washed three times with TBS-T and then imaged using a ChemiDoc MP Imaging System (BioRad).

Co-interacting protein identification technology or CoPIT developed by Pankow et al. was employed for affinity purification of CFTR bound with interactors as described previously^38^. Briefly, for immunoprecipitation, cell lysates were precleared with 4B Sepharose beads (Sigma) at 4°C for 1 hour while rocking. Precleared lysates were then immunoprecipitated with Protein G beads complexed to the 24-1 antibody (6 mg antibody per mL of beads) overnight at 4°C while rocking. The next day, a sample of the supernatant was collected for Western blot analysis as the “clear”, indicating residual protein not isolated by the beads. Resin was washed three times with TNI buffer, washed twice with TN buffer, and frozen at −80 °C for 1.5 hours. Proteins were then eluted twice with shaking at 37 °C for 20 minutes - 1 hour with a 0.2 M Glycine (pH 2.3) /0.5% IGEPAL CA-630 buffer. Samples were then blotted as described above or processed for mass spectrometry analysis.

### Multiplexed LC-MS/MS

Eluted samples were precipitated in methanol/chloroform, washed three times with methanol, and air dried. Protein pellets were then resuspended in 3 µL 1% Rapigest SF Surfactant (Waters) followed by the addition of 10 µL of 50mM HEPES pH 8.0, and 32.5µL of H_2_O. Samples were reduced with 5mM tris(2-carboxyethyl)phosphine (TCEP) (Sigma) and alkylated with 10mM iodoacetamide (Sigma). 0.5 µg of Trypsin (Sequencing Grade, Promega or Pierce) was then added and incubated for 16-18 hours at 37°C, with shaking at 700 rpm. Peptide samples were then reacted with TMT 11-plex reagents (Thermo Fisher) in 40% v/v acetonitrile and incubated for one hour at room temperature. Reactions were then quenched by the addition of ammonium bicarbonate (0.4% w/v final concentration) and incubated for one hour at room temperature. TMT labeled samples for a given experiment were then pooled and acidified with 5% formic acid (Sigma, v/v). Samples were concentrated using a SpeedVac and resuspended in buffer A (95% water, 4.9% acetonitrile, and 0.1% formic acid, v/v/v). Cleaved Rapigest SF surfactant was removed by centrifugation for 30 minutes at 21,100 x g. Peptides were directly loaded onto a triphasic MudPIT columns using a high-pressure chamber. Samples were then washed for 30 minutes with buffer A. LC-MS/MS analysis was performed using an Exploris 480 (Thermo Fisher) mass spectrometer equipped with an UltiMate3000 RSLCnano System (Thermo Fisher). MudPIT experiments were performed with 10 µL sequential injections of 0, 10, 30, 60, & 100% buffer C (500mM ammonium acetate in buffer A), followed by a final injection of 90% buffer C with 10% buffer B (99.9% acetonitrile, 0.1% formic acid v/v) and each step followed by a 90 minute gradient from 4% to 40% B with a flow rate of either 300 or 500 nL/min, followed by a 15 minute gradient from 40% to 80% B with a flow rate of 500 nL/min. on a 20cm fused silica microcapillary column (ID 100 µM) ending with a laser-pulled tip filled with Aqua C18, 3 µm, 100Å resin (Phenomenex). Electrospray ionization (ESI) was performed directly from the analytical column by applying a voltage of 2.0 or 2.2 kV with an inlet capillary temperature of 275°C. Data-dependent acquisition of MS/MS spectra was performed by scanning from 300-1800 m/z with a resolution of 60,000 to 120,000. Peptides with an intensity above 1.0E4 with charge state 2-6 from each full scan were fragmented by HCD using normalized collision energy of 35 to 38, with a 0.4 m/z isolation window, 120 ms maximum injection time, at a resolution of 15,000-45,000 scanned from 100 to 1800 m/z or defined a first mass at 110 m/z and dynamic exclusion set to 45 or 60s, and a mass tolerance of 10 ppm.

### Interactome Characterization

Peptide identification and TMT-based protein quantification was carried out using Proteome Discoverer 2.4. MS/MS spectra were extracted from Thermo XCaliber .raw file format and searched using SEQUEST against a UniProt human proteome database (released 03/25/2014) containing 20,337 protein entries. The database was curated to remove redundant protein and splice-isoformsand supplemented with common biological MS contaminants. Searches were carried out using a decoy database of reversed peptide sequences and the following parameters: 10 ppm peptide precursor tolerance, 0.02 Da fragment mass tolerance, minimum peptide length of 6 amino acids, trypsin cleavage with a maximum of two missed cleavages, static cysteine modification of 57.0215 (carbamidomethylation), and static N-terminal and lysine modifications of 229.1629 (TMT sixplex). SEQUEST search results were filtered using Percolator to minimize the peptide false discovery rate to 1% and a minimum of two peptides per protein identification. TMT reporter ion intensities were quantified using the Reporter Ion Quantification processing node in Proteome Discoverer 2.4 and summed for peptides belonging to the same protein.

### Proteostasis Modulator Experiments

Western blot analysis was performed on P67L, L206W, G85E, and F508del CFTR under a combination of VX-809 and proteostasis modulator conditions. A 10 cm dish was transiently transfected with 4 ug of the respective CFTR mutation using calcium phosphate transfection described previously. After 16 hours the media was changed and left to incubate for 1 hour. Next, the 10 cm dish was split using 1 mL trypsin and 5 mL media to inoculate a 6 well dish with 1 mL of transfected cells such that the transfection efficiency was the same across the conditions. Next, 2 μL of 3 mM VX-809 was added to respective wells of the 6-well dish for a final concentration of 3 nM. Then, each column of the 6 well dish was treated with DMSO, 10 μM bortezomib, or 10 μM E-64 respectively. Cells were collected 12-16 hours after treatment, lysed, normalized, and run for Western blotting as described above.

### Pathway Analysis/Selection of Biological Pathway from GOterms on Uniprot

First, we searched our dataset against the pathway classification presented by Pankow et al^28^. This step classified all previously identified interactors. For novel interactors we manually annotated pathways by searching gene names on UniProt and classifying pathways by GO-terms for Biological Pathway. Some pathways were simplified such as Membrane Organization which constitutes lipid raft organizing proteins, proteins involved in ER and PM organization, as well as Golgi proteins. All proteins comprising the ribosome were classified as translation. All proteins comprising the Hsp70 and Hsp90 cycle were classified or re-classified as folding (e.g. BAG2 is an Hsp70 co-chaperone that stimulates nucleotide exchange but is also involved in proteasomal degradation). Degradation on the other hand was divided into distinct categories: proteasomal degradation, autophagy, and non-specific degradation. Any proteasomal subunit, protein containing proteasome in the GO terms, or proteasomal degradation in the Function description on unitprot.org was placed in proteasomal degradation. Any protein containing terms “autophagy”, “autolysosome”, “micro-autophagy”, “ER-phagy”, “autophagosome maturation”, or “aggresome” were categorized as Autophagy. UBR4 was classified as Autophagy as it is calmodulin binding, an important autophagy protein. Any proteins involved in degradation that did not fit the former two categories were categorized as Degradation.

### Statistical Analysis

To determine statistically significant CFTR interaction proteins, we used a two-tailed paired t-test (using the scipy.stats.ttest_rel package in Python) to calculate the p-value between the log2 TMT intensity of each protein pulled down with CFTR and the corresponding the log2 TMT intensity of each protein pulled down with tdTomato control. First, we normalized log2 protein abundances to the log2 CFTR protein abundance across TMT channels in individual TMT11plex sets (**Supplemental Figure 2**). We then scaled the log2 fold abundances for these interactors to the corresponding WT DMSO condition within each run to provide a common reference point. Subsequently, we averaged these scaled log2 abundance changes from replicate TMT11plex sets. Finally, we excluded any proteins not quantified for both WT and F508del since direct quantitative comparison is not possible for these proteins. A heatmap portraying all proteins included for all pathways and all mutants showing this normalization is included (**Supplemental Figure S3B**) as well as the data plotted (**Supplement Dataset 3**). For aggregate pathways statistics in violin plots, we used a one-way ANOVA with Geisser-Greenhouse correction and post-hoc Tukey multi-comparison testing to evaluate the statistically significant difference between conditions for a given pathway. Data plotted is included (**Supplement Dataset 4**). Finally, we used a two-tailed ratio paired t-test to determine statistical differences between quantified Western blot data.

## ACKNOWLEDGEMENTS

We thank Drs. Eric Sorscher and Jeong Wong (Emory University, Atlanta, Georgia, USA) for CFTR expression plasmids. We thank members of the Plate lab for their critical reading and feedback of this manuscript. This work was funded by T32 GM065086 (NIGMS) (EFM); Cystic Fibrosis Foundation Postdoctoral Fellowhip (SABUSA19F0) (CMPS); R35 GM133552 (NIGMS); and Vanderbilt University funds.

## AUTHOR CONTRIBUTIONS

LP and CS conceived the study and planned the experiments. EM and CS performed the experiments. EM, LP, and CS analyzed the data, prepared the figures, and wrote the manuscript. MK assisted with experiments and data analysis. LP, CS, and MK critically reviewed the manuscript. LP and EM edited the manuscript.

## DATA AVAILABILITY

The mass spectrometry proteomics data have been deposited to the ProteomeXchange Consortium via the PRIDE partner repository with the dataset identifier XXXXX. All other necessary data are contained within the manuscript.

