## Supporting Information for "Distinct proteostasis states drive pharmacologic chaperone susceptibility for Cystic Fibrosis Transmembrane Conductance Regulator misfolding mutants"

for

**TABLE OF CONTENT**

Supplemental Figures S1 – S7 pp 2-11
Supporting Information References p 11

### SUPPLEMENTAL FIGURES

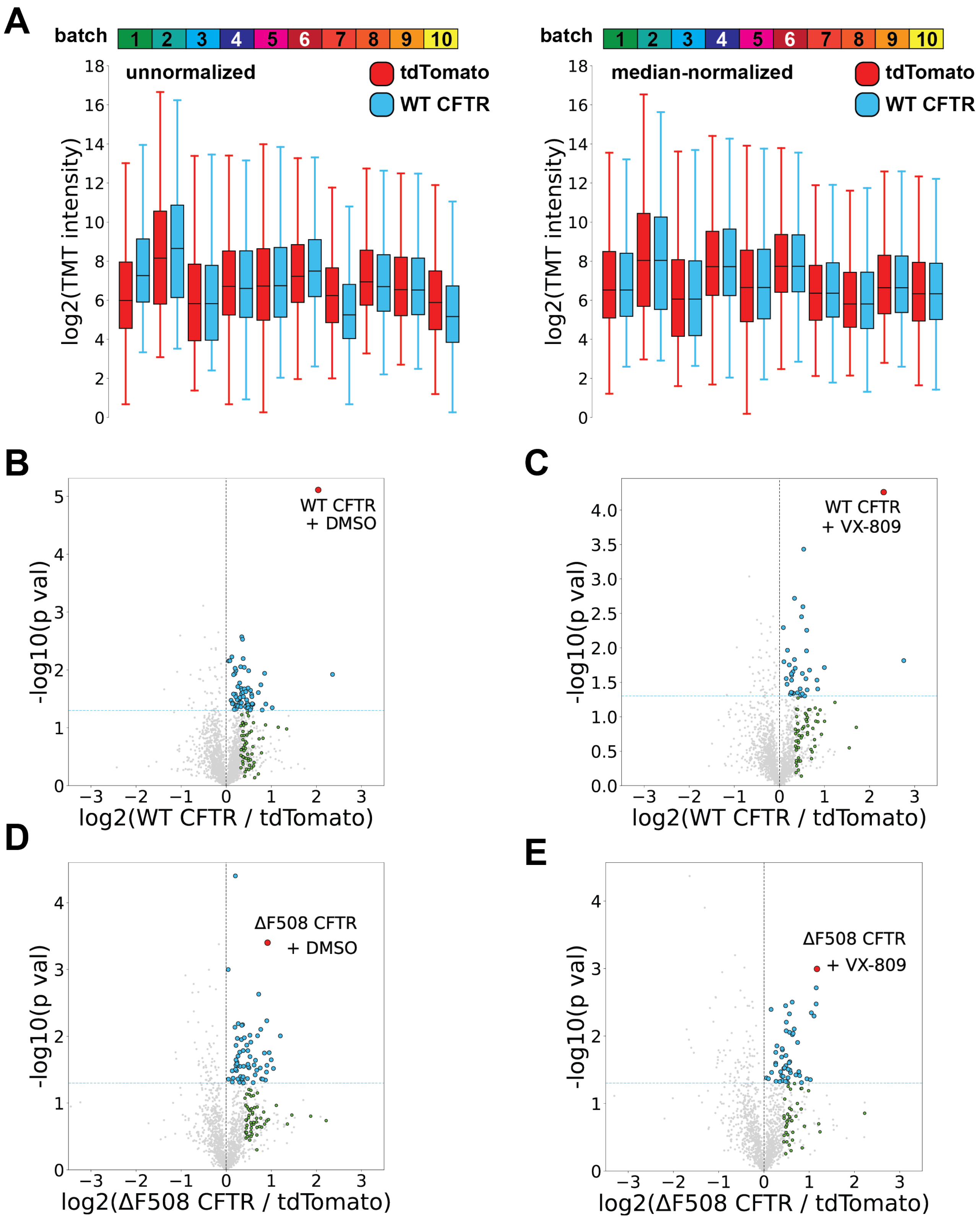


**Supplemental Figure S1. Quantification and identification of WT and F508del CFTR interactors. A.** Box and whisker plot showing the distribution of TMT intensities in the tdTomato and WT CFTR TMT channels for all 10 TMT labeled LC-MS/MS runs used in this study. Median normalization of all TMT channels across a given run makes the median in each run for various conditions approximately equal, as represented by tdTomato control and WT CFTR with DMSO. The box limits represent the upper and lower quartile with a line at the median, the whiskers represent the limits 1.5 times the interquartile range. **B-E.** Individual volcano plots for identification of WT and F508del interactors with DMSO or VX-809 treatment using multiplexed AP-MS proteomics. The x axis represents log 2-fold change of CFTR IP over IP of tdTomato mock control. A statistical significance cutoff of $p < 0.05$ is portrayed by a blue dotted line. Blue dots represent newly identified statistically significant interactors in this study and green dots represent previously identified interactors[1,2] that did not meet our statistical cutoff threshold but were enriched greater than one standard deviation among all log2(FC) between CFTR vs. tdTomato control. **B.** WT CFTR with DMSO treatment. **C.** WT CFTR with 3 μM VX-809 treatment. **D.** F508del CFTR with DMSO treatment **E.** F508del CFTR with 3 μM VX-809 treatment.

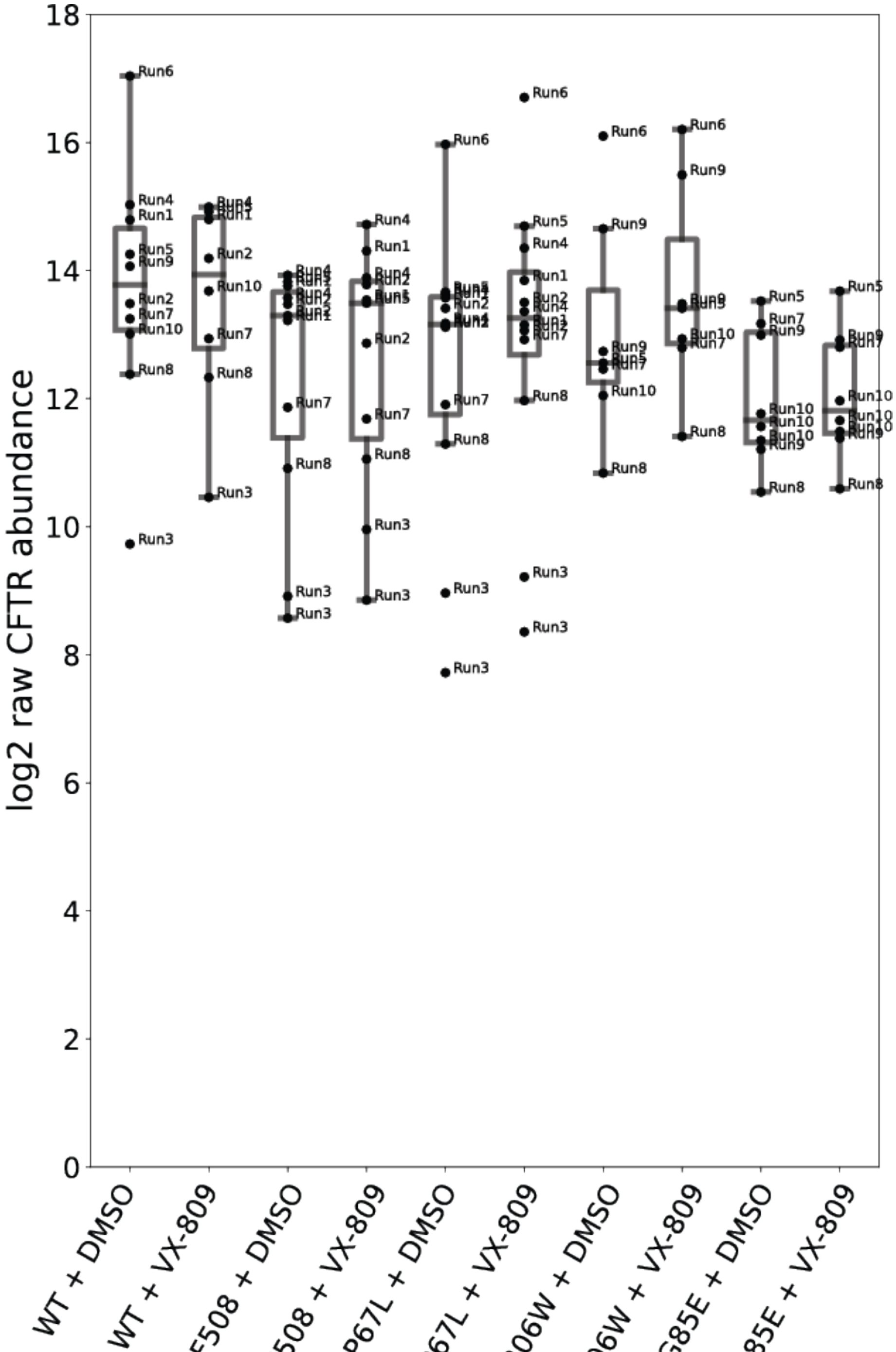


**Supplemental Figure S2. Quantification of CFTR levels across all TMT runs demonstrates lower CFTR expression of mutants compared to WT.** Log2 TMT intensity of quantified CFTR of each variant in each run. Individual CFTR quantifications are plotted as labeled dots, and box and whisker plot represent the distribution of CFTR quantification for a particular mutant. The box limits represent the upper and lower quartile with a line at the median, the whiskers represent 1.5 times the interquartile range. This plot reveals higher TMT intensity for WT CFTR than most mutants, as well as some variation in CFTR levels for the mutants. Due to the variability of CFTR quantification across samples normalizing interactors to the CFTR level in the specific TMT channel is required for direct quantitative comparison.

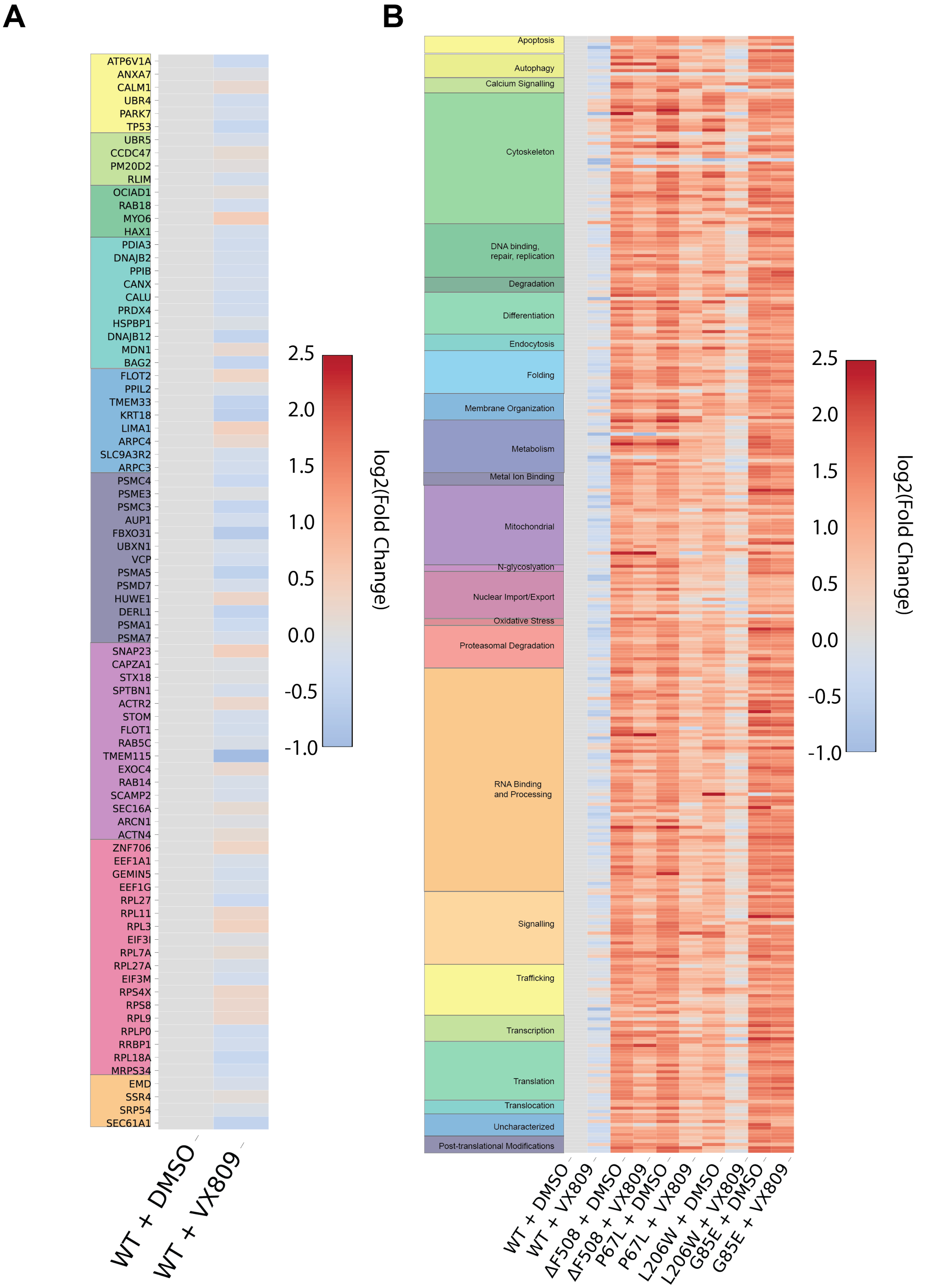


**Supplemental Figure S3. Heatmaps of WT interactors under VX-809 treatment and all variants with and without VX-809 reveals the alterations to CFTR protein interactions occurring with VX-809. A.** A pathway ordered heatmap comparing the interactions of WT CFTR treated with 3 μM VX-809 scaled to WT with DMSO. Proteins were selected from manually curated list of proteostasis interactors and pathways were assigned with GO-terms. **B.** A pathway order heatmap comparing WT, F508del, P67L, L206W, and G85E interactions under DMSO and VX-809 conditions for all pathways annotated in our master list. P67L and L206W demonstrate a reduction in interactions across pathways including those not directly involved in proteostasis.

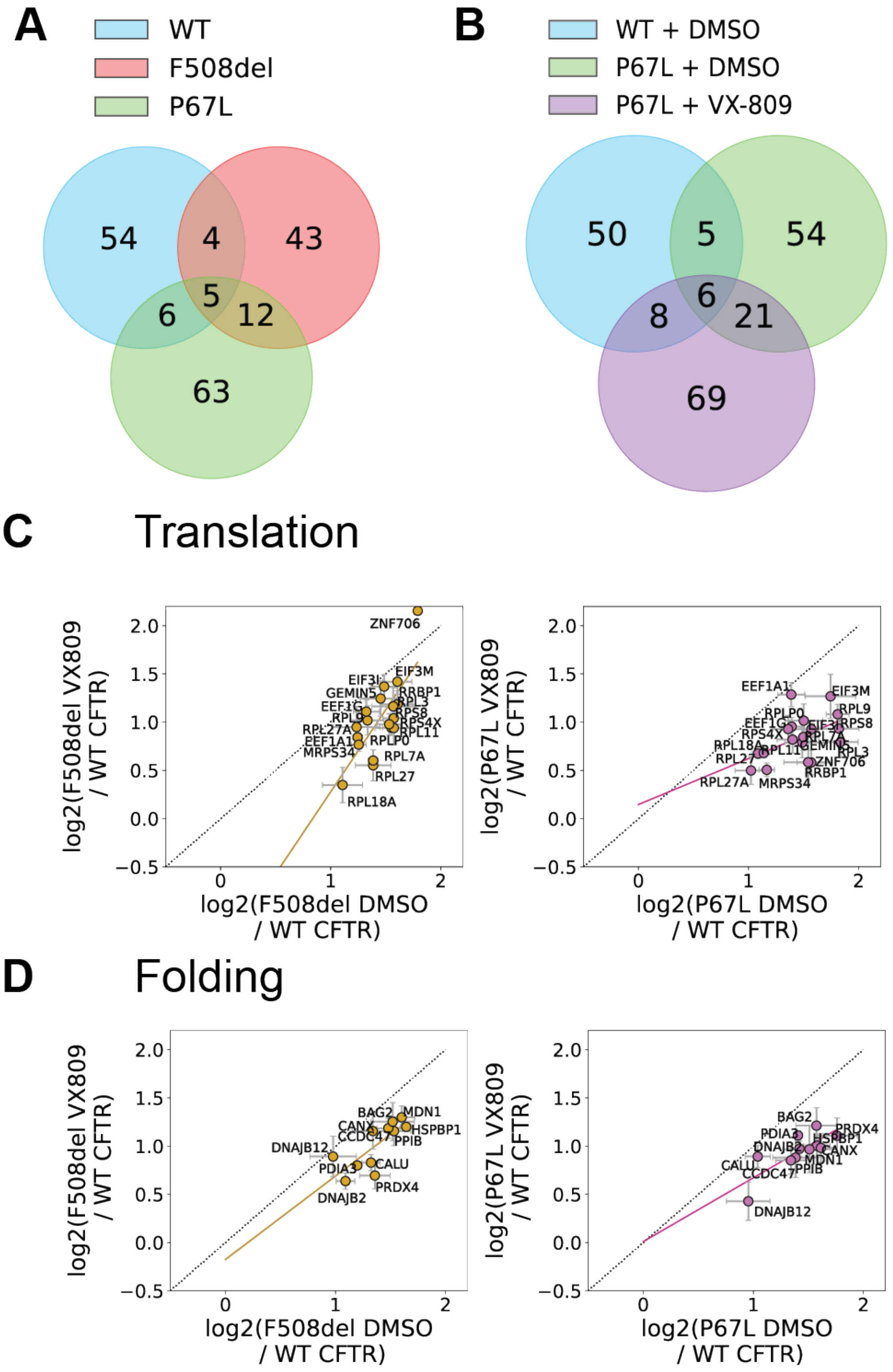


**Supplemental Figure S4. P67L and F508del exhibit similar protein interactions indicative of their characterization as class-II misfolding variant. A.** Overlap in statistically significant identified interactors for WT CFTR treated with DMSO, F508del CFTR with DMSO, and P67L CFTR with DMSO. P67L and F508del share more interactors with one another than they share with WT, consistent with the notion that class-II CFTR variants experience similar interactomes. **B.** Overlap in statistically significant interactors between WT CFTR with DMSO, P67L with DMSO, and P67L with 3 μM VX-809. P67L shares more interactors with WT when treated with VX-809 consistent with the notion that the interactomes can be remodeled under favorably rescuing conditions as suggested by Pankow et al.[2] **C.** Translational protein quantifications correlated for F508del and P67L CFTR between DMSO and 3 μM VX-809 treatment normalized to WT with DMSO. Black dotted line represents a normal line with a slope of 1 and intercept at the origin. The gold and purple dotted line is the linear least squared best fit of the translational proteins for F508del and P67L interaction quantification respectively. **D** Folding protein quantifications correlated for F508del and P67L CFTR between DMSO and 3 μM VX-809 treatment normalized to WT with DMSO. Black dotted line represents a normal line with a slope of 1 and intercept at the origin. The gold and purple dotted line is the linear least squared best fit of the folding proteins for F508del and P67L interaction quantification respectively.

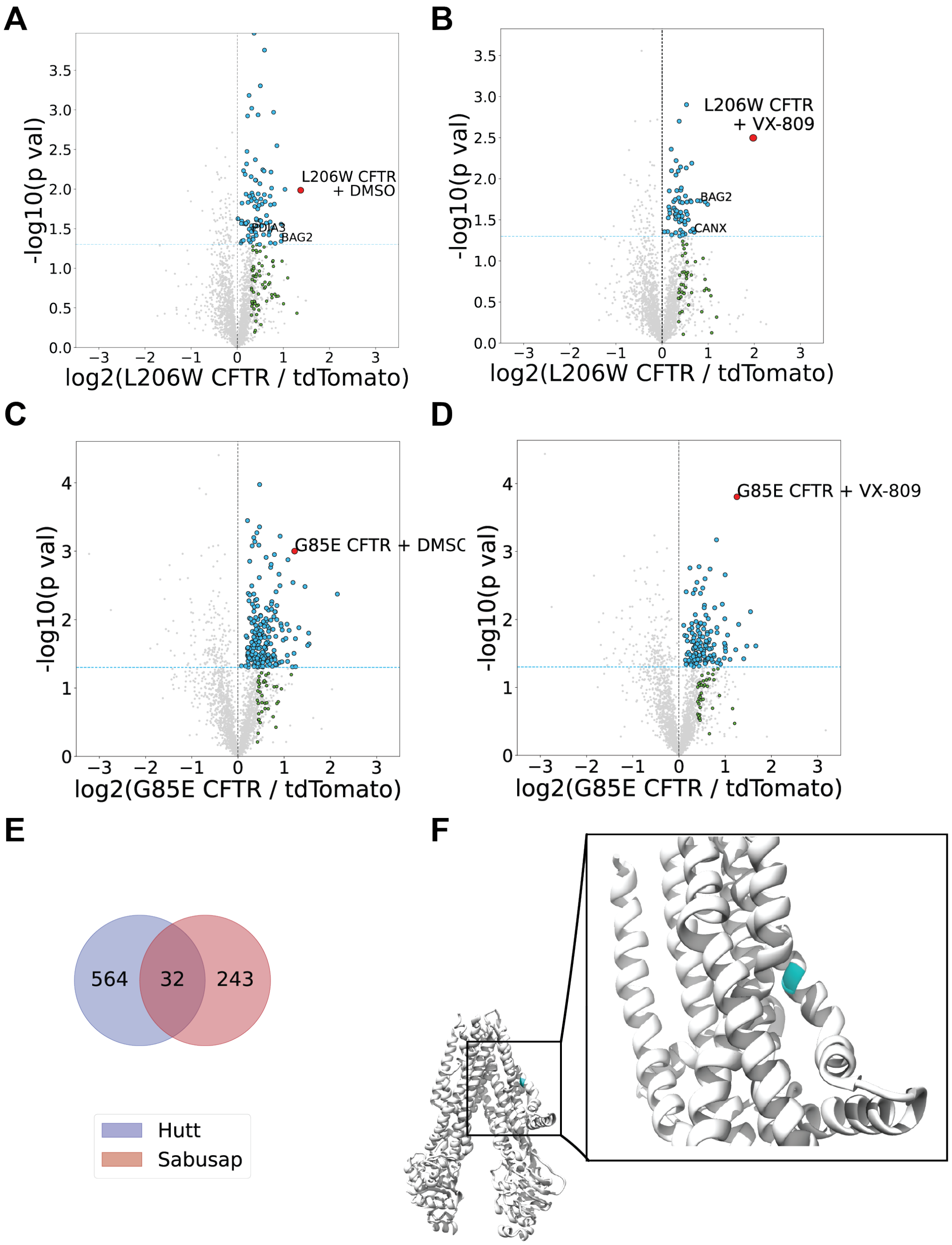


**Supplemental Figure S5. Identification of statistically significant interactors of L206W CFTR and G85E CFTR with and without VX-809 treatment. A-D.** Volcano plots for the identification of L206W and G85E interactors using multiplexed AP-MS proteomics. The x axis represents log 2-fold change of CFTR IP over IP of tdTomato mock control. A statistical significance cutoff of $p < 0.05$ is portrayed by a blue dotted line. Blue dots represent newly identified statistically significant interactors in this study and green dots represent previously identified interactors[1,2] that did not meet our statistical cutoff threshold but were greater than one standard deviation among all log2(FC) between CFTR vs. tdTomato control. **A.** Identification of novel L206W CFTR interactors with DMSO treatment. **B.** Identification of L206W CFTR interactors with 3 μM VX-809 treatment **C.** Identification of G85E CFTR interactors with DMSO treatment. **D.** Identification of G85E CFTR interactors with 3 μM VX-809 treatment. **E.** Overlap of interactors for G85E CFTR with DMSO newly identified in this study with those interactor identified by Hutt, et al[1]. **F.** G85, shown in blue, is in the first transmembrane domain helix of TMD1. Shown here is the closed state CFTR structure (PDB ID 5UAK)[3]. Introduction of a large, charged residue at this location likely hinders insertion or stability in the membrane.

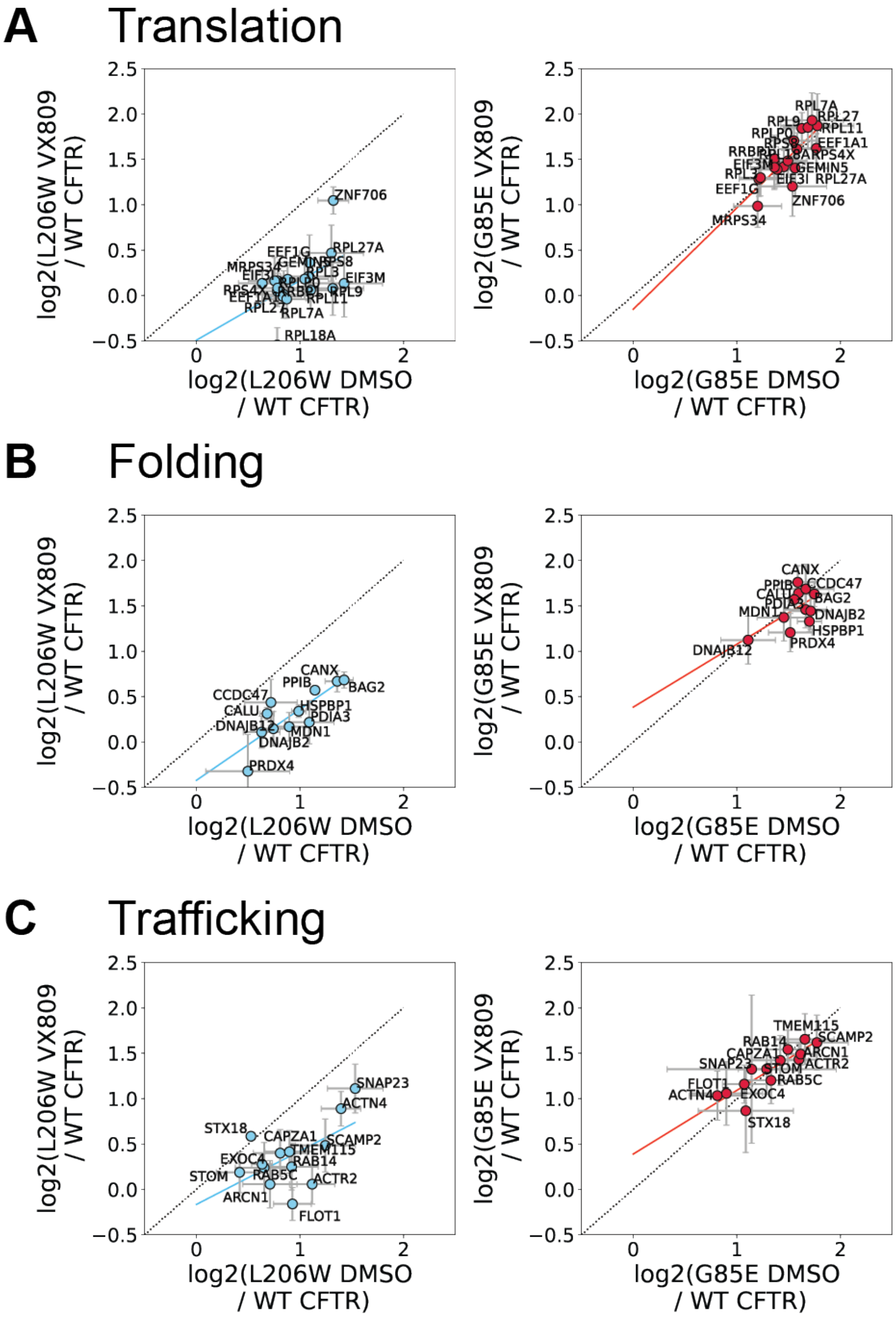


**Supplemental Figure S6. VX-809 alters protein interactions levels for L206W, moving linear best fit below the normal line compared to G85E. A.** Translational protein quantifications correlated for L206W and G85E CFTR between DMSO and 3 μM VX-809 treatment normalized to WT with DMSO. Black dotted line represents a normal line with a slope of 1 and intercept at the origin. The blue and red dotted line is the linear least squared best fit of the translational proteins for L206W and G85E interaction quantification respectively. **B.** Folding protein quantifications correlated for L206W and G85E CFTR between DMSO and 3 μM VX-809 treatment normalized to WT with DMSO. Black dotted line represents a normal line with a slope of 1 and intercept at the origin. The blue and red dotted line is the linear least squared best fit of the folding proteins for L206W and G85E CFTR interaction quantification respectively. **C.** Trafficking protein quantifications correlated for L206W and G85E CFTR between DMSO and 3 μM VX-809 treatment normalized to WT with DMSO. Black dotted line represents a normal line with a slope of 1 and intercept at the origin. The blue and red dotted line is the linear least squared best fit of the trafficking proteins for L206W and G85E interaction quantification respectively.

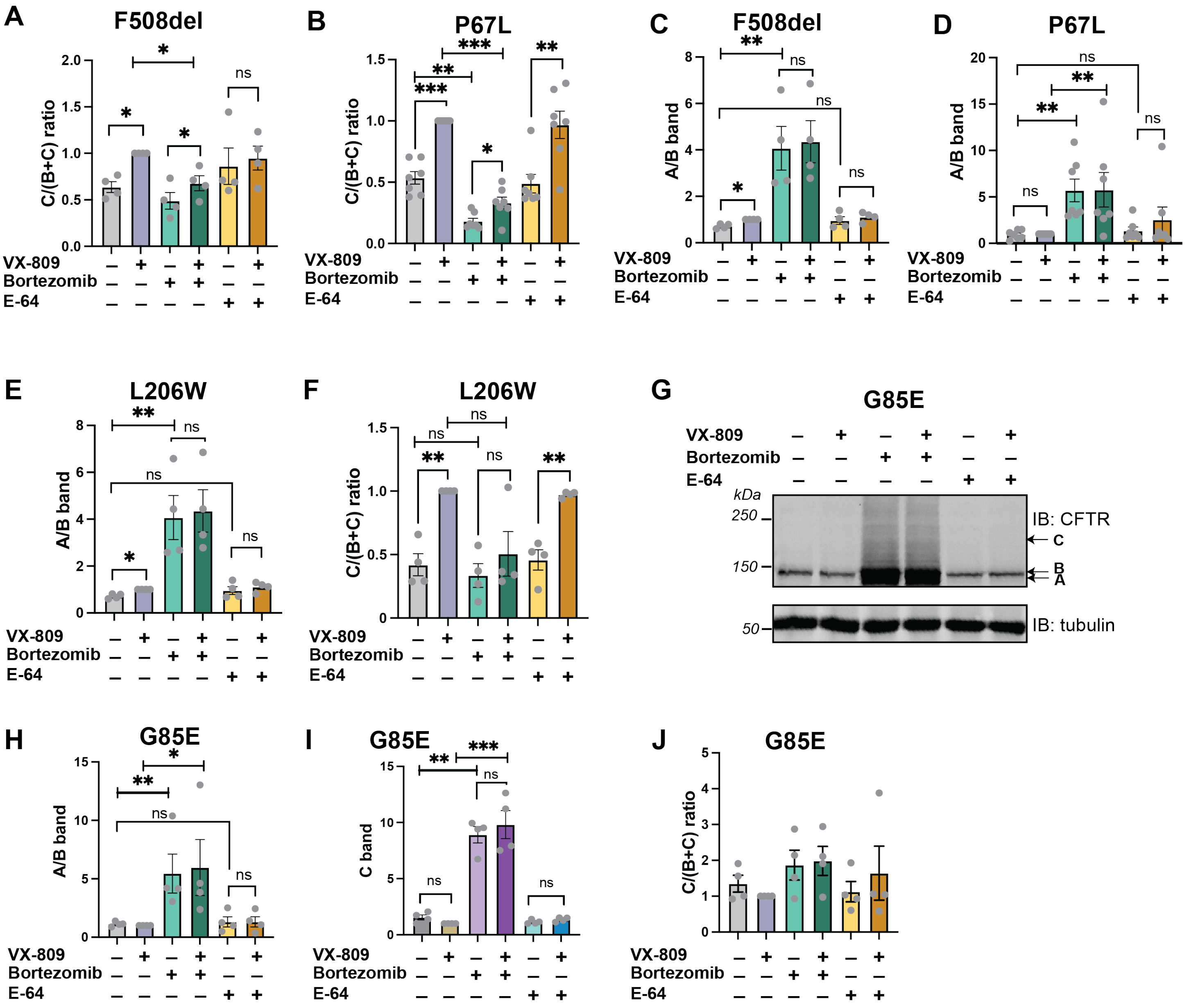


**Supplemental Figure S7. Altering proteostasis with proteasomal and autophagy inhibition uniquely changes the VX-809 response of CFTR mutants.** **A.** Quantification of F508del CFTR trafficking efficiency (C band over C band + B band) Western blot data showing the relative ratio of trafficked to total F508del CFTR normalized to the 3 µM VX-809 treatment condition. Representative Western blot is shown in **Figure 6A**. Individual measurements are shown as grey points, error bars represent standard error of the mean (n=4), statistical significance calculated with a paired, two tailed student t-test for all plots and p values depicted by * < 0.05, ** < 0.01, *** < 0.001, and **** < 0.0001. **B.** Quantification of P67L CFTR trafficking efficiency (C band over C band + B band) from Western blot data showing the relative ratio of trafficked to total P67L CFTR normalized to the 3 µM VX-809 treatment condition. Representative Western blot is shown in **Figure 6B. C.** Quantification of immature (A/B band) F508del with 3 µM VX-809, 10 µM bortezomib, 10 µM E-64, or combination of treatments western blot data. These bar graphs show the relative ER-retained F508del CFTR normalized to the 3 µM VX-809 treatment condition. **D.** Quantification of immature (A/B band) P67L with 3 µM VX-809, 10 µM bortezomib, 10 µM E-64, or combination of treatments western blot data. These bar graphs show the relative ER-retained P67L CFTR normalized to the 3 µM VX-809 treatment condition. **E.** Quantification of immature (A/B band) L206W with 3 µM VX-809, 10 µM bortezomib, 10 µM E-64, or combination of treatments western blot data. These bar graphs show the relative ER retained P67L CFTR normalized to the 3 µM VX-809 treatment condition. **F.** Quantification of L206W CFTR trafficking efficiency (C band over C band + B band) from Western blot data showing the relative ratio of trafficked to total L206W CFTR normalized to the 3 μM VX-809 treatment condition. Representative Western blot is shown in **Figure 6I**. **G.** Representative Western blot of G85E CFTR trafficking assay under 3 μM VX-809, 10 μM bortezomib, 10 μM E-64, and combined treatments. Blots were probed with 217 CFTR antibody and tubulin antibody as a loading control. **H.** Quantification of immature (A/B band) G85E with 3 μM VX-809, 10 μM bortezomib, 10 μM E-64, or combination of treatments from Western blot data in **G**. These bar graphs show the relative ER retained F508del CFTR normalized to the 3 μM VX-809 treatment condition. **I.** Quantification of mature (C band) G85E from Western blot data in **G.** showing the relative post-Golgi G85E CFTR normalized to the 3 μM VX-809 treatment condition. **J.** Quantification of G85E CFTR trafficking efficiency (C band over C band + B band) from western blot in **G.** data showing the relative ratio of trafficked to total G85E CFTR normalized to the 3 μM VX-809 treatment condition.
